# „APP family member dimeric complexes are formed predominantly in synaptic compartments“

**DOI:** 10.1101/2022.09.13.507759

**Authors:** S. Schilling, A. August, M. Meleux, C. Conradt, L. Tremmel, S. Teigler, V. Adam, E.H. Koo, S. Kins, S. Eggert

## Abstract

The amyloid precursor protein (APP), a key player in Alzheime’s disease (AD), is part of a larger gene family, including the APP like proteins APLP1 and APLP2. They share similar structures, form homo- and heterotypic dimers and exhibit overlapping functions. We investigated complex formation of the APP family members via two inducible dimerization systems, the FKBP-rapamycin based dimerization as well as cysteine induced dimerization, combined with coimmunoprecipitations and Blue Native (BN) gel analyses. Within the APP family, APLP1 shows the highest degree of dimerization and high molecular weight (HMW) complex formation. Interestingly, about 20% of APP is dimerized in cultured cells while about 50% of APP is dimerized in mouse brains, independent of age and splice forms. Furthermore, we could show that dimerized APP originates mostly from neurons and is enriched in synaptosomes. Finally, BN gel analysis of human cortex samples shows a significant decrease of APP dimers in AD patients compared to controls, suggesting that loss of dimers of full-length APP might correlate with loss of synapses in the process of AD.

## INTRODUCTION

The amyloid precursor protein (APP) plays an essential role in Alzheimer’s disease, since sequential cleavages of β- and γ-secretase lead to the formation of the 4 kDa Aβ peptide, which accumulates in brains of AD patients (Hardy and Selkoe, 2002). APP is part of a larger gene family, which includes two mammalian homologues, the amyloid precursor like protein 1 and 2 (APLP1 and APLP2) (Muller and Zheng, 2012). The APP family members share several common features like neuronal localization including prefrontal cortex and processing by α-, β-, and γ-secretase (Eggert et al., 2004; Li and Sudhof, 2004; Lorent et al., 1995; Mallm et al., 2010; Muller and Zheng, 2012). Combined and single gene mutations of the APP family show impairment in spatial learning, long-term potentiation (Ring et al., 2007; Weyer et al., 2014), decreased number of dendritic spines and profound neuromuscular junction (NMJ) defects (Schilling et al., 2017; Tyan et al., 2012; Wang et al., 2009; Weyer et al., 2014). Thus, loss of the physiological function of APP/APLPs might contribute to loss of synapses in AD (Scheff et al., 1990). This is underlined by two studies of differently generated conditional APP/APLP1/APLP2 triple KO mice. Selective inactivation of the APP family members in excitatory neurons of the postnatal forebrain via CamK2a-Cre recombination resulted in impaired hippocampal synaptic plasticity, learning, and memory as well as neuronal hyperexcitability (Lee et al., 2020). This was indicated by increased neuronal spiking via Kv7 channels in response to depolarizing current injections. In contrast, neither cortical neurodegeneration nor increases in apoptosis and gliosis up to 2 years of age have been found (Lee et al., 2020). A different study analyzing conditional APP/APLP1/APLP2 triple KO (cTKO) mice lacking the APP family in excitatory forebrain neurons due to Nex-Cre recombination, revealed disrupted hippocampal lamination, impaired long-term potentiation, reduced dendritic length and spine density (Steubler et al., 2021).

APP/APLPs also share a common domain structure (Coulson et al., 2000). Their large ectodomains include the so called E1 and E2 domains (Muller and Zheng, 2012). It has been shown that APP, APLP1, and APLP2 can dimerize in a homotypic and heterotypic manner (Schilling et al., 2017; Soba et al., 2005; Wang et al., 2009). Thereby, the E1 domain has been identified as the main interface of *cis*-as well as *trans*-cellular dimerization (Baumkotter et al., 2012; Kaden et al., 2009; Soba et al., 2005). So far, different physiological functions have been implied for *cis*- and *trans*- dimerization of APP/APLPs. It was shown that *cis*-homodimerization of APP (Eggert et al., 2009) as well as *cis*-heterodimerization of APP with the APP homologues APLP1 or APLP2 (Kaden et al., 2009) decreases Aβ production. APP *cis*-dimerization leads to an altered localization presumably via interaction changes with LRP1 and differences in retrograde transport with SorLA, two known risk factors in AD (Eggert et al., 2017; Kang et al., 2000; Schmidt et al., 2017). In line with those results, a report from Willnow and colleagues showed a strong increase of APP dimers in brains of SorLA KO mice, underlining an interplay of the AD risc factor SorLA and APP dimerization (Schmidt et al., 2012).

Significantly, *trans*-dimerization of APP proteins has been reported to promote cell-cell adhesion (Schilling et al., 2017; Soba et al., 2005; Stahl et al., 2014; Wang et al., 2009). This is supporting the notion that APP and its homologues might function as synaptic cell adhesion proteins (Schilling et al., 2017). Therefore, we aimed to investigate APP /APLP dimerization in brain in more detail, in this study.

## MATERIAL AND METHODS

### Plasmids

The following plasmids were used: Generation of APP F1 pC4 F1 has been described (Eggert et al., 2009). For cloning of APLP1 pC4 F1, the vector pC4 F1 (Clontech) has been digested EcoRI – XbaI. APLP1 WT in pBluescript SK+ was digested with EcoRI and BamHI to obtain a 1358 bp fragment. A PCR was performed to add an XbaI site at the 3’ end of the APLP1 ORF. APLP1 WT in pBluescript Sk+ served as a template (Paliga et al., 1997). The sense primer starts 30-50 bp before the BamHI site in the APLP1 ORF 5‘ GAGCAGAAGGAACAGAGGCA 3’ and the antisense primer including XbaI was as follows 5’ GTCAGTTCTAGAGGGTCGTTCCTCCAGGAAG 3’. The PCR product was digested BamHI and XbaI and ligated with the EcoRI – BamHI fragment via EcoRI and XbaI in vector pC4 F1. APLP2 pC4 F1 was cloned by cutting dimerization vector pC4 F1 (Clontech) EcoRI – XbaI. APLP2-763 WT in pCEP4 (Eggert et al., 2004) was digested EcoRI – XhoI to obtain a 2100 bp fragment. A PCR was performed using sense primer 5’ C ATG GTC ATT GAC GAG ACT C 3’ starting ∼ 30 bp before the internal unique XhoI site of APLP2-763 ORF. The following antisense primer 5’ GTCAGCTCTAGAAATCTGCATCTGCTCCAGG 3’ was used to append an XbaI site to the 3’ end of the APLP2 ORF. The PCR product was cut XhoI – XbaI. The EcoRI – XhoI fragment and the XhoI – XbaI fragment were ligated EcoRI – XbaI in dimerization vector pC4 F1.

Generation of APLP1 CT HA pcDNA3.1+ neo has been described (Schilling et al., 2017). Cysteine dimerization mutants APLP1 R579C CT HA pcDNA3.1+ neo, APLP1 E580C CT HA pcDNA3.1+ neo were cloned via site directed mutagenesis and verified by sequencing.

Cloning of APLP2 CT HA pcDNA3.1+ neo has been described (Soba et al., 2005). Cysteine dimerization mutants APLP2 CT HA L690C pcDNA3.1+ neo, APLP2 CT HA S692C pcDNA3.1+ neo were obtained via site directed mutagenesis and verified by sequencing.

Generation of APP695 CT HA pcDNA3.1+neo has been described (Eggert et al., 2009). To obtain APP751 CT HA pcDNA3.1+neo, an HA tag has been appended to the C-terminus of APP751 WT pcDNA3.1 + neo via PCR. Sense primer 5’ GAAGTTGAGCCTGTTGATGCC 3’ starts at position 1848 in the hAPP751 ORF. Antisense primer 5’ GCTGACCTCGAGTTATGCGTAGCTGGTACGTCGTACGGATAGTTCTGCATCTGC TC 3’ contains an XhoI site and an HA tag. The 400 bp PCR product was cut BglII – XhoI to obtain a 350 bp fragment. APP751 WT pcDNA3.1 was digested EcoRI – BglII to obtain the remaining insert of 1940 bp. The vector pcDNA3.1 + neo containing the 3’ UTR of APP (Eggert et al., 2009) was cut EcoRI and XhoI to ligate in the EcoRI – BglII and BglII – XhoI fragments.

APP770 CT HA pcDNA3.1+neo was generated by appending a C-terminal HA tag to APP770 WT in pcDNA3.1 + neo via PCR. Therefore, the following sense primer was used 5’ GAAGTTGAGCCTGTTGATGCC 3’ starting at position 1906 in the hAPP770 ORF and an antisense primer 5’ GCTGACCTCGAGTTATGCGTAGTCTGGTACGTCGTACGGATAGTTCTGCATCTG CTC 3’ including an XhoI site and an HA tag. The resulting PCR product: was digested BglII – XhoI to obtain a 350 bp fragment. APP751 WT in pcDNA3.1 + was cut EcoRI – BglII to obtain the remaining insert of 1994 bp. Vector 3’ UTR APP in pcDNA3.1 + neo (Eggert et al., 2009) was digested EcoRI and XhoI. The EcoRI – BglII and BglII – XhoI fragments were ligated in the EcoRI – XhoI digested vector.

Generation of the plasmids Myc APPΔCT 648 pcDNA3.1+neo and HA APPΔCT 648 pcDNA3.1+neo was based on site directed mutagenesis by introducing a stop codon after position 648 at the C-terminal end of the APP transmembrane domain of spliceform APP695. The following primer pair was used: 5’ C ACC TTG GTG ATG CTG TGA AAG AAG AAA CAG TAC 3’ and 5’ GTA CTG TTT CTT CTT TCA CAG CAT CAC CAA GGT G 3’. N-terminally c-myc tagged APLP1ΔCT and c-myc tagged APLP2ΔCT constructs have already been described (Soba et al., 2005). N-terminally HA tagged APLP1ΔCT pcDNA3.1+neo and N-terminally HA APLP2ΔCT pcDNA3.1+neo constructs have been generated by introducing a stop codon C-terminal to the transmembrane domain of APLP1, or APLP2, respectively.

### Antibodies

Primary rat monoclonal antibodies included anti-HA antibody (000000011867431001, 3F10, Roche, Rotkreuz, Switzerland). Primary mouse monoclonal antibodies anti-c-myc antibody (9E10) (Ab32, Abcam, Cambridge, UK), PSD-95 (Abcam, Cambridge, UK), Synaptophysin (Sigma-Aldrich), Golgi Marker GM130 (BD Biosciences). Further, the following rabbit polyclonal antibodies were used: anti-c-myc (A-14, sc-769, Santa Cruz, Nunningen, Switzerland), Y188 (APP C-terminal antibody, Epitomics), anti-APLP1 (57; (Eggert et al., 2004)), D2-II, APLP2 C-terminal antibody (Calbiochem).

### Cell lines and transfections

N2a mouse neuroblastoma cells were maintained in the following media: Minimum Essential Medium (MEM, Gibco) supplemented with 10% FBS (Sigma), 1% Penicillin/Streptomycin (Gibco), 1% L-Glutamine (200 mM) (Sigma), 1% non-essential amino acids, 1% sodium pyruvate. HeLa Kyoto cells were maintained in the following media: Dulbecco’s modified Eagle’s medium (DMEM, Gibco) supplemented with 10% FBS (Sigma), 1% L-Glutamine (200 mM) (Sigma), and 1% Penicillin/ Streptomycin (Gibco). For Western blot analysis, cells were transfected with jetPRIME (Polyplus, Illkirch, France) according to manufacturer’s instructions. The dimerizer, AP20187 (B/B) (Clontech, Saint-Germain-en-Laye France) was added 4 h post-transfection overnight.

### Mice

The generation and genotyping of knock-out lines were described previously [APP KO and APLP1 KO (Heber et al., 2000); APLP2 KO (von Koch et al., 1997). All mice have been backcrossed at least six times to C57BL/6 mice. The sex of the species used is of either sex. C57BL/6J mice [embryonic day 14 (E14)] were used for the generation of primary cortical neuron cultures. The used transgenic J20 mice express the APP Indiana (V717F) and Swedish (K670M) mutations on a BL/6 background (Mucke et al., 2000a). Mice were treated in accordance with the German law for conducting animal experiments and followed the National Institutes of Health *Guide for the Care and Use of Laboratory Animals*. Animal housing, breeding, and the sacrifice of mice were approved by the German administration. All experimental protocols were performed in accordance with the European Communities Council Directive of 24 November 1986 (86/609/EEC).

### Blue Native Gel Analysis

Blue Native gel electrophoresis was performed according to a protocol modified from (Schagger et al., 1994).

### For analysis of transiently transfected N2a cells

In brief, cells in one 10 cm cell culture dish were washed once and collected in phosphate-buffered saline at 4°C. The cell pellets were resuspended in 1 ml of homogenization buffer (250 mM sucrose in 20 mM HEPES, pH 7.4, with protease inhibitor mix “Complete”, Roche Rotkreuz, Switzerland) and then sheared by passing through a 27x gauge needle 10 times.

### For analysis of mouse brains

Mouse cortices were homogenized in homogenization buffer (250 mM sucrose in 20 mM HEPES, pH 7.4, with protease inhibitor mix “Complete”, Roche Rotkreuz, Switzerland) at 800 rpm 13 times with a glass-Teflon homogenizer on ice. The following steps are identical for cells as well mouse cortices. The Post nuclear supernatant was collected after a low-speed spin at 300 x *g* for 15 min at 4°C. The membranes were pelleted after centrifugation at 100.000 x *g* for 1 h at 4°C and washed once with 200 μl of homogenization buffer. After repeating the ultracentrifugation step, the pellets containing the membranes were resuspended in 200 μl of homogenization buffer.

50 μg of protein were solubilized with Blue Native sample buffer (1.5 M amino caproic acid, 0.05 M Bis-Tris, 10% *n*-dodecyl-β-D-maltoside, and protease inhibitor at pH 7.5). The samples were incubated on ice for 30 min and then centrifuged for 10 min at 14.000 rpm at 4°C in a microcentrifuge. Blue Native loading buffer (5.0% Serva Coomassie Brilliant Blue G250 and 1.0 M aminocaproic acid) was added to the supernatant. The samples were separated on 4–15% Tris-HCl gels (Criterion, Bio-Rad) overnight at 4 °C with Coomassie Blue containing cathode buffer (10 x cathode buffer, pH 7.0, 0.5 M Tricine, 0.15 M Bis-Tris, 0.2% Coomassie Blue) and anode buffer (pH, 0.5 M Bis-Tris). The gel was transferred to a polyvinylidene difluoride membrane. The following molecular weight standards were used: thyroglobulin (669 kDa), apoferritin (443 kDa), catalase (240 kDa), aldolase (158 kDa), and bovine serum albumin (66 kDa), all from Sigma-Aldrich, Munich, Germany.

### *Cis*-coimmunoprecipitation

*Cis*-coimmunoprecipitations were performed, as described before (Eggert et al., 2017). Briefly, N2a cells co-expressing APP CT HA and APP CT myc, APLP1 CT HA and APLP1 CT myc or APLP2 CT HA and APLP2 CT myc were harvested and lysed in 50 mM Tris/HCl pH 7.5; 150 mM NaCl; 5 mM EDTA; 1% NP40; 1:25 protease inhibitor (Complete (with EDTA), Roche, Rotkreuz, Switzerland) 18 h post-transfection. Equal volumes of cell lysates containing ∼1000 μg protein were precleared with 10 μl Protein A Sepharose (GE Healthcare, Freiburg, Germany) for 1 h at 4°C. Afterwards, the supernatant was incubated at 4°C overnight with 20 μl anti-HA antibody coated beads (Roche, Rotkreuz, Switzerland) at RT. After several washing steps with lysis buffer and 10 mM Tris/HCl pH 7.5, the beads were resuspended in SDS sample buffer (0.125 M Tris/HCl pH 6.8; 20% glycerol; 4% SDS; 0.01% bromophenol blue; 100 mM DTT) and incubated for 5 min at 95°C. The samples were loaded on an 8% Tris/glycine gel and subjected to Western blot analysis using different primary antibodies (anti-c-myc (A-14, sc-769, Santa Cruz, Nunningen, Switzerland and HA antibody, rat, 3F10, Sigma-Aldrich, Munich, Germany).

### *Trans*-coimmunoprecipitation

*Trans*-coimmunoprecipitations were performed by transiently transfecting N2a cells (10 cm dishes) using JetPrime (Polyplus) expressing APP CT HA, APP CT myc, APLP1 CT HA, APLP1 CT myc, APLP2 CT HA or APLP2 CT myc. 4 h after the transfection, the cells of following two dishes were combined in one 6 cm dish: APP CT HA and APP CT myc, APLP1 HA and APLP1 CT myc, APLP2 HA and APLP2 CT myc expressing cells. 18 h post transfection, the cells were harvested and lysed in 50 mM Tris/HCl pH 7.5; 150 mM NaCl; 5 mM EDTA; 1% NP40; 1:25 protease inhibitor (Complete (with EDTA), Roche, Rotkreuz, Switzerland). Further steps of the immunoprecipitation followed as described above for the *cis*-coimmunoprecipitation.

### Sample preparation for SDS Gels and Western blot

Cells were lysed for 15 min at 4°C in lysis buffer (50 mM Tris/HCl, pH 7.5, 150 mM NaCl, 5 mM EDTA, and 1% Nonidet P40) supplemented with protease inhibitors (CompleteTM protease inhibitor mixture, Roche, Rotkreuz, Switzerland). The supernatants were collected, and the protein concentration determined with a BCA assay (Sigma, Deisenhofen, Germany). Equal amounts of protein samples were separated on 8% Tris/glycine gels and then transferred to nitrocellulose membranes (GE Healthcare, Uppsala, Sweden). Subsequently the membranes were incubated with primary antibodies and HRP-coupled secondary antibodies (Jackson ImmunoResearch, West Grove, Pennsylvania, USA). Chemiluminescence was measured, using an imager and the software Fusion (Vilber Loumat).

### Synaptosomal preparation and enrichment of the postsynaptic density

Samples were always kept on ice, and all centrifugation steps were performed at 4°C. One mouse brain was homogenized in solution A (0.32 M sucrose; 1 mM NaHCO3; 1 mM MgCl2; 0.5 mM CaCl2) by a Potter S Homogenizer. Centrifugation for 10 min at 800 x *g* sedimented crude cell fragments [“post-nuclear supernatant” (PNF)]. The supernatant was centrifuged for 15 min at 9000 x *g* (smaller cell components stay in the supernatant). The pellet was resuspended in solution A, centrifuged again for 15 min at 10,000 x *g*, and resuspended in solution B (0.32 M sucrose; 1 mM NaHCO3) to obtain a raw synaptosomal fraction (Syn raw). Synaptosomal membranes were isolated via hypo osmotic shock by the addition of double-distilled water. The reaction was stopped with 0.5 M HEPES/NaOH, pH 7.4 [fraction “synaptosomes after hypoosmotic shock” (Syn Hyp)]. The solution was centrifuged for 20 min at 25,000 x *g*, and the pellet was subsequently (dissolved in solution B) loaded on a sucrose gradient (0.5 M - 1 M - 2 M sucrose in 1 mM NaHCO_3_). A discontinuous density centrifugation at 82,500 x *g* was performed for 3 h. The low dense fraction (Syn L) and the high dense fraction were collected (Syn HdFr) and centrifuged again at 201,800 x g for 20 min. The Pellets were subsequently resuspended in 5 mM Tris/HCl (PSD L and PSD H). To analyze equal amounts of proteins, a BCA test was performed, and the samples were loaded on an 8% Tris/glycine gel or a Blue Native gel.

### Immunocytochemistry

HeLa cells were seeded at a density of 35,000 cells per 24-well plate (Greiner) on 14-mm coverslips and transfected via jetPrime. The cells were fixed after 18-20 h for 10 min at 37°C in 4% PFA with 4% sucrose and permeabilized for 10 min with 0.1% NP40. After incubation of primary antibodies at 4°C overnight and secondary antibodies for 1 h at RT (Alexa Flour 488 and 594) cells were embedded in Mowiol (Sigma-Aldrich) and subjected to imaging with the software Axiovision 4.8 at the microscope Axio Observer Z.1.

### Cell surface staining

HeLa cells were seeded at a density of 35,000 cells per 24-well plate (Greiner) on 14-mm coverslips and transfected via jetPrime. After 18-20 h the cells were cooled on ice and incubated with an α-c-myc antibody to stain only proteins which were localized at the surface. Then the cells were fixed for 10 min on ice in 4% PFA with 4% sucrose and another 20 min at RT. Then they were incubated with the secondary antibody Alexa Flour 488 for 1 h at RT. After washing the cells were permeabilized for 10 min with 0.1% NP40 and stained again with the same myc antibody to also visualize the intracellular proteins. The secondary antibody Alexa Flour 594 was again incubated for 1 h at RT. Then the cells were embedded in Mowiol (Sigma-Aldrich) and subjected to imaging with the software Axiovision 4.8 at the microscope Axio Observer Z.1.

## RESULTS

We investigated complex formation of all APP family members via an inducible dimerization system which we already established for APP (Eggert et al., 2018a; Eggert et al., 2009). A 12 kDa FKBP tag adjacent to an HA tag was fused to the C-terminus of APP695, APLP1, and APLP2 763, termed APP F1, APLP1 F1, and APLP2 F1, respectively **(Figure 1A)**. Western Blot analysis of heterologously expressed fusion proteins in N2a cells, which had been lysed in buffer containing 1% NP40, revealed for APP F1, APLP1 F1, and APLP2 F1 an apparent molecular weight of ∼130 kDa, 100 kDa and 140 kDa, as expected **(Figure 1B)**.

**Figure 1:**
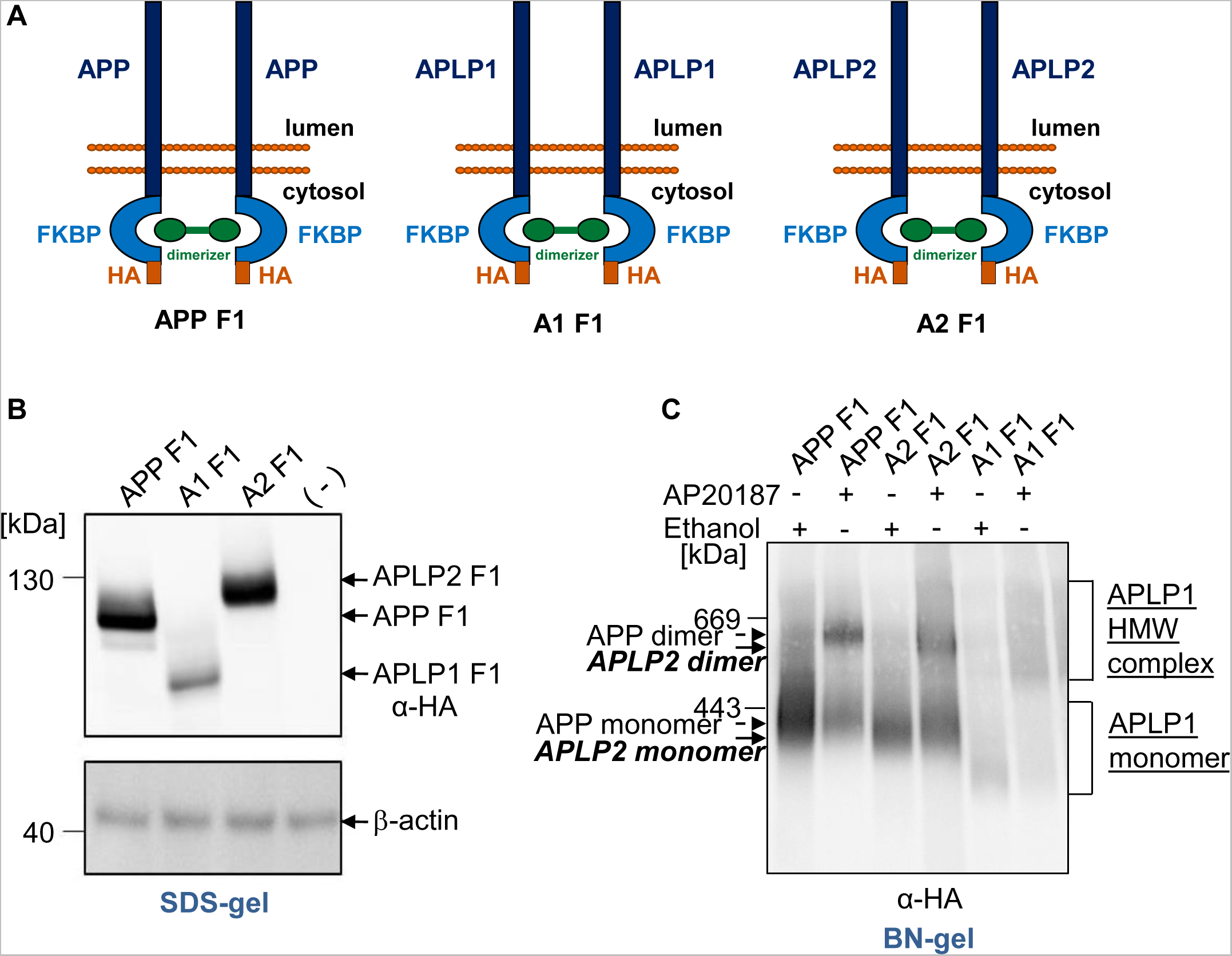
Comparative analysis of FKBP induced dimerization of the APP gene family members on denaturing SDS gels and semi-denaturing Blue Native gels. **(A)** Schematic illustration of the used constructs APP F1, APLP1 F1 or APLP2 F1. By addition of the rapamycin analogue AP20187 the FKBP tags can be dimerized. **(B)** The FKBP tagged APP gene family members were heterologously expressed in N2a cells. Non-transfected N2a cells served as a negative control. Equal amounts of cell lysates were separated on SDS-Gels and Western blot detection followed with an α-HA antibody. Detection with primary antibody β-actin was performed as a loading control. **(C)** APP F1, APLP1 F1 or APLP2 F1 were heterologously expressed in N2a cells and treated with 100 nM of the rapamycin analogue AP20187 overnight to induce dimerization of APP/APLPs. Incubation with the solvent of AP20187, ethanol, served as a negative control. The samples were analyzed via Blue Native PAGE and Western blot analysis followed with an α-HA antibody.

Addition of an analogue of rapamycin to transiently transfected N2a cells, which is cell permeable, will allow efficient dimerization of our proteins of interest, the APP family members, via binding of two FKBP molecules (Clackson, 2006; Eggert et al., 2018a; Eggert et al., 2009). N2a cells heterologously expressing APP F1, APLP1 F1 or APLP2 F1 were either treated with the vehicle ethanol as a control or with 100 nM of AP20180, the so called dimerizer, overnight. The cells were harvested in homogenization buffer without detergent and disrupted mechanically. Then, the complexes were solubilized out of the membrane using the mild detergent N-dodecyl-β-D-maltoside and separated on a gradient gel to detect complexes of APP, APLP1 or APLP2. For APP and APLP2, we visualized two distinct signals at 600 – 700 kDa and ∼200 - 300 kDa **(Figure 1C)**. Similar to APP (Eggert et al., 2018a; Eggert et al., 2009), for APLP2, mainly two distinct signals with >80% in the low molecular weight (LMW) complex and only a minor part, <20%, in the high molecular weight (HMW) complex were detected in ethanol treated control cells. In contrast, APLP2 extracted from dimerizer treated cells showed about >60% of APLP2 in the HMW complex. Due to the almost identical apparent molecular weight compared with APP, these data indicate that the HMW signals of APLP2 represent dimeric APLP2, as observed for APP before (Eggert et al., 2018a; Eggert et al., 2009). In contrast, APLP1 shows a different pattern compared to APP and APLP2 in the ethanol treated control with signals between 200 and 800 kDa, possibly representing monomeric and dimeric complexes between 200 - 400 kDa and 400 - 800 kDa, respectively **(Figure 1C)**.

To corroborate our assumption that the HMW complexes represent dimeric forms of APLPs, we analyzed cysteine induced APLP1 and APLP2 dimers, analogously to cysteine induced dimerization of APP, which has already been described using APP L17C and APP K28C containing amino acid substitutions within the Aβ domain (Eggert et al., 2018a; Eggert et al., 2009; Munter et al., 2007; Scheuermann et al., 2001). For this purpose, we were introducing cysteines N-terminally to the transmembrane domain of APLP1 and APLP2, assumed to cause formation of disulfide bridges between the two molecules containing the cysteine variants. We generated APLP1 containing the amino acid changes R597C and E580C **(Figure 2A)** as well as the APLP2 variants L690C and S692C **(Figure 2B)** using N-terminally HA tagged constructs and firstly examined APLP1 R597C and E580C via SDS gel electrophoresis compared to APLP1 WT. Non transfected N2a cells served as a negative control. We prepared cell lysates with or without DTT, allowing visualization of APLP1 covalent dimers on SDS gels, since disulfide bridges will remain intact. Western blot detection with an α-HA antibody revealed that for APLP1 R597C, a higher amount of APLP1 dimers was visualized compared to APLP1 E580C while for APLP1 WT almost no SDS resistant APLP1 dimers were present **(Figure 2C)**. This shows that APLP1 dimerization can be induced by incorporation of cysteines in their juxtamembraneous domains. Furthermore, these data indicate that dimer formation via covalent binding does not occur for WT APLP1.

**Figure 2:**
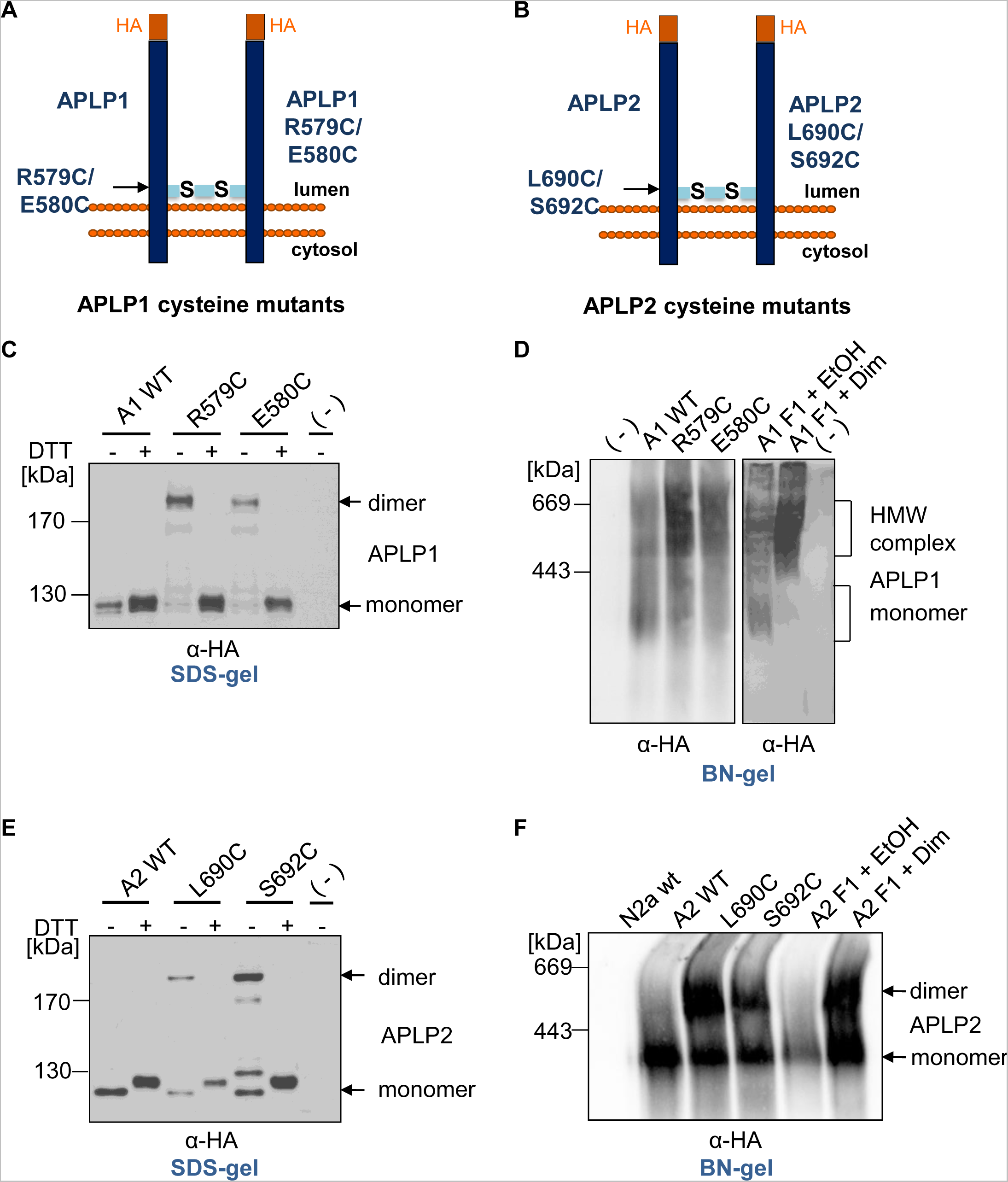
Comparative analysis of induced dimerization of APLP1 and APLP2 with cysteine mutants on denaturing SDS gels and semi-denaturing Blue Native gels. **(A/B)** Schematic illustration of the used cysteine mutants for APLP1 and APLP2 **(C)** APLP1 WT, APLP1 R579C and APLP1 E580C were heterologously expressed in N2a cells. Non-transfected N2a cells served as a negative control. Equal amounts of cell lysates were prepared in samples with or without DTT, to allow visualization of cysteine dimers via disulfide bridges. Separation on SDS-gels followed and Western blot detection with an α-HA antibody. **(D)** APLP1 WT, APLP1 R579C, APLP1 E580C, and APLP1 F1 were heterologously expressed in N2a cells. APLP1 F1 expressing cells were either treated with 100 nM of the rapamycin analogue AP20187 overnight to induce dimerization of APLP1 or with the solvent ethanol as a negative control. Non transfected N2a cells served as a further negative control. The samples were analyzed via Blue Native PAGE and Western blot analysis followed with an α-HA antibody. **(E)** APLP2 WT, APLP2 L690C, and APLP2 S692C were transiently transfected in N2a cells. Equal amounts of protein were denatured in sample buffer with and without DTT and analyzed after SDS PAGE via Western blot detection with α-HA antibody. **(F)** APLP2 WT, APLP2 L690C, APLP2 S692C, and APLP2 F1 were heterologously expressed in N2a cells. APLP2 F1 expressing cells were either treated with 100 nM of the rapamycin analogue AP20187 overnight to induce dimerization of APLP2 or with the same volume of the vehicle ethanol as a negative control. Non transfected N2a cells served as a further negative control. The samples were analyzed via Blue Native PAGE and Western blot analysis followed with an α-HA antibody.

We aimed to investigate high molecular weight complexes of APLP1 via BN gel analyses. In order to assign the detected signals correctly to APLP1 monomers and dimers on BN gels, we compared induced dimer formation of APLP1 via cysteine mutants with induced dimer formation via the FKBP-rapamycin system on BN gels which revealed that APLP1 R597C and E580C show an accumulation of APLP1 high molecular weight complexes at about the same apparent molecular weight between 400 - 800 kDa as APLP1 F1 transfected cells treated with dimerizer overnight **(Figure 2D)**. Induced dimer formation was >50% for both systems.

We performed the same kind of investigation for APLP2 and firstly prepared cell lysates of N2a cells expressing either APLP2 WT, APLP2 L690C or APLP2 S692C. SDS gel analysis followed by Western blot detection with an α-HA antibody revealed strong dimer formation for APLP2 S692C as well as APLP2 L690C in samples prepared using sample buffer without DTT. APLP2 WT did not show dimer formation under these conditions indicating that dimerization of APLP2 WT is not based on covalent interactions similarly as shown before for APLP1 WT **(Figure 2C, E)**. Next, we examined, analogously to APLP1, induced dimer formation of APLP2 via cysteine mutants in comparison to induced dimer formation via the FKBP-rapamycin system on BN gels. This indicated that APLP2 L690C and S692C show an accumulation of APLP2 dimers at the same apparent molecular weight as APLP2 F1 transfected cells treated with dimerizer overnight at ∼600 kDa and monomeric APLP2 at ∼300 kDa **(Figure 2F)**. Induction of APLP2 dimer formation was ∼66% for both systems. Similarly, to APP WT and APP F1, APLP2 WT and APLP2 F1 showed only very little dimer formation of about 16% **(Figure 1C, 2F)**.

Since we observed a markedly higher rate of APLP1 dimerization compared to APP and APLP2 via BN gel analysis, even without using a dimerization system, we wanted to confirm this result via a different method. Therefore, we were choosing to analyze the dimerization state of the APP family members via co-immunoprecipitation **(Figure 3A)**. N2a cells heterologously expressing myc and HA tagged APP, APLP1 or APLP2, respectively, were lysed 24 h post-transfection and subjected for co-immunoprecipitation with anti-HA antibody coupled beads. N2a cells co-transfected with vector and APP HA served as a control. Western blot detection with an α-c-myc antibody revealed comparable expression level of the c-myc tagged APP, APLP1, and APLP2 proteins with the lowest amount for APLP1 **(Figure 3B)**. In contrast, Western blot detection of the IP samples with an α-c-myc antibody demonstrated for APLP1 the strongest interaction of the APP family members, while APLP2 indicated the weakest homodimerization, which was confirmed by quantification of the integrated densities **(Figure 3C)**. Western blot detection of the same membrane with an α-HA antibody showed comparable signals of HA tagged APP/APLPs in the input control as well as of the immunoprecipitated proteins **(Figure 3B)**. Notably, the myc and HA tagged APP/APLP proteins were expressed in the same cells and thus the observed interaction likely took place in a lateral *cis*-directed fashion within the membrane. However, the interaction might also partially take place between APP/APLPs of neighbouring cells in a *trans*-cellular fashion.

**Figure 3:**
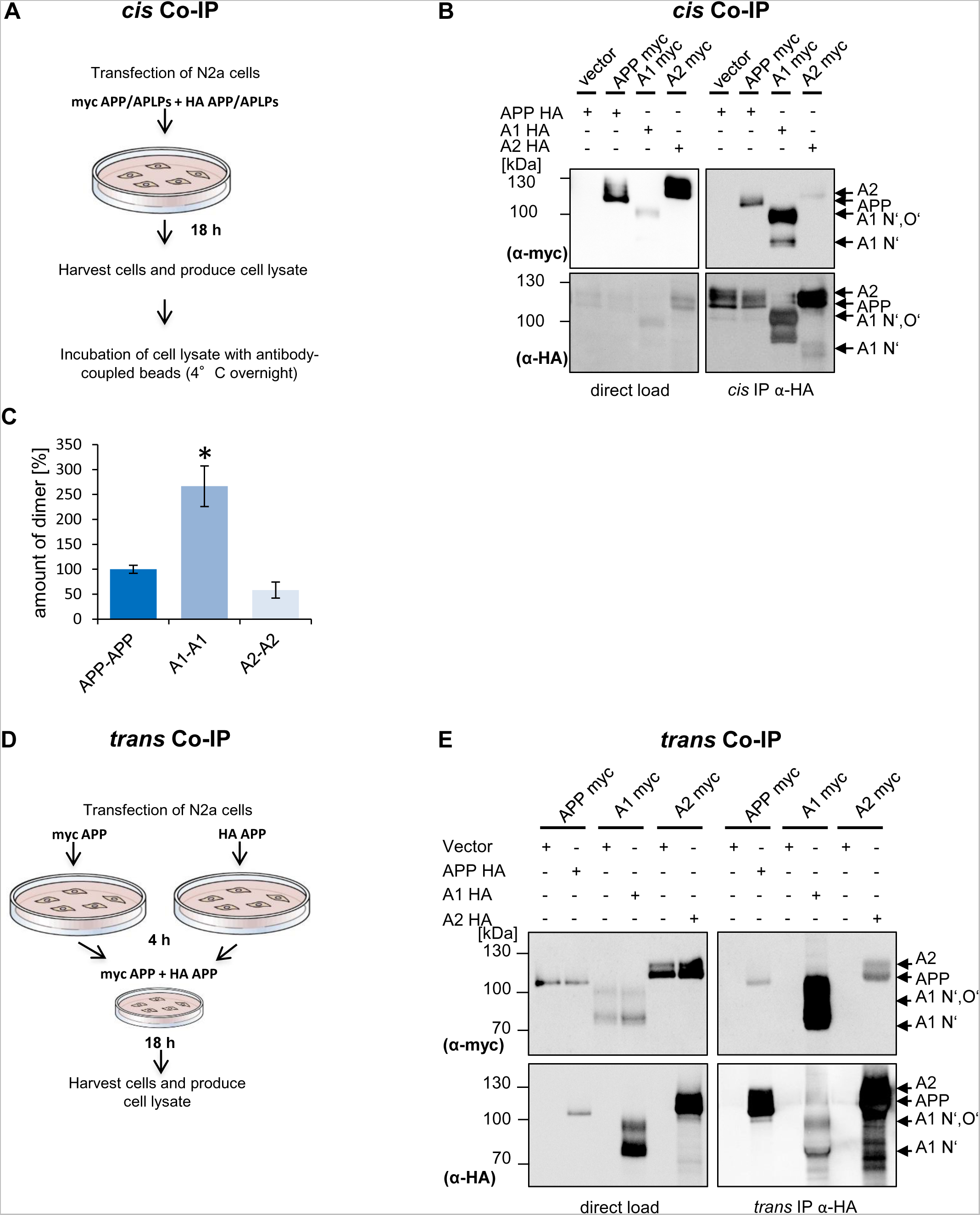
The extent of APP, APLP1, APLP2 *cis*-dimers and *trans*-dimers in N2a cells. **(A)** Experimental design of the *cis***-**coimmunoprecipitation. **(B)** *Cis***-** coimmunoprecipitation of APP, APLP1, and APLP2 (*cis*-homodimers). N2a cells were transiently co-transfected with APP myc and APP HA, APLP1 myc and APLP1 HA or APLP2 myc and APLP2 HA constructs. Cotransfection of APP HA and empty vector served as a negative control. Equal amounts of cell lysates were loaded directly on an SDS gel and analyzed via Western blot with primary α-HA or α-myc antibodies (input controls). Further, equal amounts of cell lysates were used for immunoprecipitation with α-HA antibody coated agarose beads. The samples were separated on an SDS gel and subjected for Western blot detection with the primary antibody α-c-myc to detect the coimmunoprecipitated proteins The same membrane was incubated afterwards with α-HA antibody to detect total amounts of immunoprecipitated APP HA, APLP1 HA or APLP2 HA. **(C)** Quantification of data shown in A. *Bars* represent mean values ± SEM; *n* = 3, unpaired Student’s *t* test \**p* < 0.05, \*\**p* < 0.01, \*\*\**p* < 0.001. **(D)** Experimental design of the *trans***-**coimmunoprecipitation. **(E)** *trans*-coimmunoprecipitation of APP, APLP1 or APLP2 *trans*-homodimers. N2a cells were transiently transfected in separate dishes with APP myc, APP HA, APLP1 myc, APLP1 HA, APLP2 myc or APLP2 HA constructs. Further, transfections of APP myc or empty vector served as a negative control. 4 h post-transfection, the following transfected cells of separate dishes were combined: APP myc and APP HA, APLP1 myc and APLP1 HA, APLP2 myc and APLP2 HA. Equal amounts of cell lysates were loaded directly on an SDS gel and analyzed via Western blot with primary α-HA or α-myc antibodies (input controls). Further, equal amounts of cell lysates were used for immunoprecipitation with α-HA antibody coated agarose beads. The samples were separated on an SDS gel and subjected for Western blot detection with the primary antibody α-c-myc to detect the coimmunoprecipitated proteins. The same membrane was incubated afterwards with α-HA antibody to detect total amounts of immunoprecipitated APP HA, APLP1 HA or APLP2 HA.

To test more directly for *trans*-interaction of the APP family members, we modified the co-immunoprecipitation assay. N2a cells expressing either myc tagged APP or HA tagged APP were grown on separate cell dishes for one day post-transfection and then co-cultivated on one plate for one more day allowing formation of *trans*-cellular contacts. **(Figure 3D)**. A small part of the cell lysates was loaded as an input control and the major part was used for the immunoprecipitation with HA beads. Western blot detection with an α-c-myc antibody revealed clearly that c-myc-tagged APP can be co-immunoprecipitated with HA-tagged APP expressed in the neighbouring cells, indicating that APP can form *trans*-cellular dimers. Combination of N2a cells transfected with vector and c-myc tagged APP served as negative control.

The same experiments were also carried out with N2a cells expressing HA or c-myc tagged APLP1 or APLP2 on separate cell plates that were again co-cultivated for one day, lysed and subjected for co-immunoprecipitation. Combination of N2a cells transfected with vector and c-myc tagged APLP1 or APLP2 served again as negative controls. Interestingly, the highest amount of *trans*-interaction was detected for APLP1 **(Figure 3E)**. Taken together, we were able two show via *cis*-as well as via *trans-* coimmunoprecipitation that APLP1 has a higher dimerization rate in comparison to its homologues APP and APLP2.

Next, we addressed the question if the stronger presence of APLP1 *trans-*dimers is maybe based on higher APLP1 cell surface levels compared to the other APP family members. All APP family members are known to be internalized via clathrin mediated endocytosis (Eggert et al., 2009; Schilling et al., 2017) with the lowest internalization rate for APLP1, which therefore shows the highest presence at the cell surface (Schilling et al., 2017). To validate that a higher cell surface localization of APP will also result in a higher amount of *trans*-dimers, we performed *trans*-coimmunoprecipitations of APPΔCT lacking the entire C-terminus and therefore all endocytosis motifs and compared it with APP WT **(Figure 4A)**. N-terminally HA tagged and N-terminally c-myc tagged APPΔCT constructs or N-terminally HA tagged and N-terminally c-myc tagged APP WT were transiently transfected into N2a cells. The differently tagged APP WT or APPΔCT expressing cells were combined 4 h after transfection and analyzed the following day. The HA tagged proteins were pulled down via HA beads and equal amounts of protein were loaded as input controls. Western blot detection of the input controls and IP samples with an anti-HA antibody confirmed that equal amounts of protein were loaded and that equal amounts of APP WT and APPΔCT were pulled down with HA beads **(Figure 4B)**. In contrast, detection with anti-c-myc antibodies demonstrated that higher amounts of APPΔCT were pulled down via *trans*-coimmunoprecipitation compared with APP WT **(Figure 4B)**. This demonstrates that indeed, a higher presence of APP at the cell surface due to strongly impaired endocytosis (Eggert et al., 2020) goes along with a higher amount of APP *trans*-dimers.

**Figure 4:**
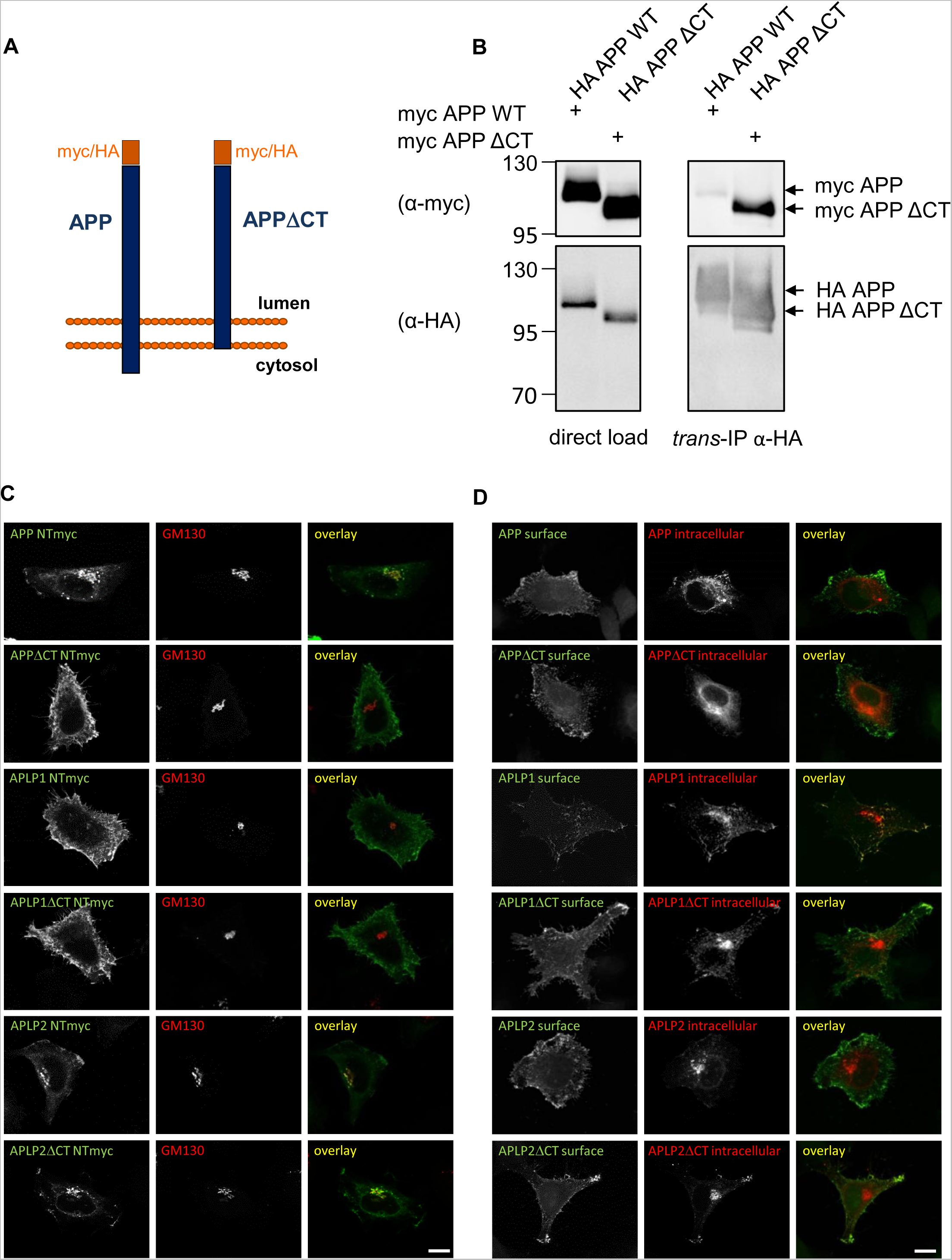
The APP gene family members lacking the C-terminus are localized to a higher amount to the cell surface which leads presumably to higher *trans*-dimerization. **(A)** Schematic representation of the N-terminally HA/myc tagged APP WT and APPΔCT constructs used. **(B)** *Trans*-coimmunoprecipitation of APP WT or APPΔCT, *trans*-homodimers. N2a cells were transiently transfected in separate dishes with myc APP, HA APP, myc APPΔCT or HA APPΔCT constructs. 4 h post-transfection, the following cells of separate dishes were combined: myc APP and HA APP, myc APPΔCT and HA APPΔCT. Equal amounts of cell lysates were loaded directly on an SDS gel and analyzed via Western blot with primary α-HA or α-c-myc antibodies (input controls). Further, equal amounts of cell lysates were used for immunoprecipitation with α-HA antibody coated agarose beads. The samples were separated on an SDS gel and subjected for Western blot detection with the primary antibody α-c-myc to detect the coimmunoprecipitated proteins. The same membrane was incubated afterwards with an α-HA antibody to detect total amounts of immunoprecipitated HA APP HA or HA APPΔCT. **(C)** HeLa cells were transiently transfected with N-terminally c-myc tagged APP, APPΔCT, APLP1, APLP1ΔCT, APLP2 and APLP2ΔCT. The cells were stained with anti-c-myc antibody to visualize the overexpressed proteins and GM130 antibody to show the *cis*-Golgi apparatus in permeabilized cells. The APP gene family members lacking the C-terminus are located to a higher amount to the cell surface. Scale bar, 10 µm. **(D)** Cell surface staining of HeLa cells transiently transfected with N-terminally c-myc tagged APP, APPΔCT, APLP1, APLP1ΔCT, APLP2 or APLP2ΔCT. Cells were incubated with an α-c-myc antibody on ice to stain only proteins which were localized at the surface. After fixation, the cells were permeabilized and stained again with an α-c-myc antibody to also visualize the intracellular proteins.

To further validate the impact of the C-terminus on the presence at the cell surface for all APP family members, we performed an immunocytochemical staining with N-terminally c-myc tagged full length proteins and mutants lacking the entire C-terminus of all APP family members (APP WT, APLP1 WT, APLP2 WT, APPΔCT, APLP1ΔCT, and APLP2ΔCT). Firstly, we were analyzing localization of the APP family members in cells permeabilized with 0.1% NP40 using an anti-c-myc antibody and the *cis*-Golgi marker GM130 **(Figure 4C)**. APP WT showed a prominent Golgi and a vesicular staining with very little signal at the cell surface, while APPΔCT seemed almost exclusively to be present at the plasma membrane. For APLP1 WT and APLP1ΔCT, mostly cell surface staining and almost no co-localization with the *cis*-Golgi marker was observed. APLP1 WT showed also a punctate cytosolic localization, which was not present for APLP1ΔCT **(Figure 4C)**. For APLP2 WT, a substantial presence in the *cis*-Golgi apparatus has been visualized with almost no additional punctate staining, but a considerable presence at the cell surface, which was quite comparable to the staining pattern of APLP2ΔCT **(Figure 4C)**. This suggests that lack of the APLP2 C-terminus including the internalization motifs surprisingly did not lead to a major shift to the presence at the cell surface.

Secondly, we were analyzing localization of the APP family members via cell surface (non-permeabilized cells) and subsequent intracellular (permeabilized cells) staining using an α-c-myc antibody combined with two different secondary antibodies (surface: Alexa Fluor-488, *green*; intracellular: Alexa Fluor-594, *red*). The overlay indicates relative intensities of total (*red*) and cell surface (*green*) APP/APLPs **(Figure 4D)**. This set of immunocytochemical stainings clearly confirms that all APP family members are present at the cell surface. Taken together, we demonstrated in cell culture cells that APLP1 shows the highest dimerization rate of the APP family members as well as the strongest presence at the cell surface.

As a next step, we wanted to analyze complex formation of the APP family members in intact tissue. Therefore, we were comparing the FKBP fusion proteins of the APP family members heterologously expressed in N2a cells after induced dimerization with WT cortical mouse brain samples via BN gel analysis **(Figure 5)**. The corresponding KO brain samples served as a control to prove specificity of the antibodies used. APP F1 expressing cells treated with the vehicle control ethanol showed about 22.17% ±4.7% APP dimers, as expected and ∼65.8% ±8.2% induced APP dimers. In contrast, APP dimers in WT mouse brains were detected with an amount of ∼48.9% ±1.6%, a ∼2x higher value compared to APP dimers in neuronal N2a cells **(Figure 5A, B)**. For APLP1 F1, about 31.8% ±6.1% APLP1 dimers were detected in ethanol treated N2a cells while induced dimerization resulted in 55.8% ±6.1% APLP1 dimers. In contrast, APLP1 in cortical mouse brains was dimerized 45.6% ±3.9%, similarly as APP **(Figure 5C, D)**. APLP2 F1 expressing cells treated with the vehicle control ethanol showed about 16.24% ±3.4% APLP2 dimers and 66.15% ±4.3% induced APLP2 dimers, similarly as for APP F1. In contrast, less APLP2 dimers were identified in WT mouse brains compared with APP and APLP1 with an amount of 26.07% ±3.60% **(Figure 5E, F)**. Taken together, dimerization of APP in mouse brains was strongly increased compared to cell culture N2a cells while dimerization of APLP1 and APLP2 was comparable between the two systems.

**Figure 5:**
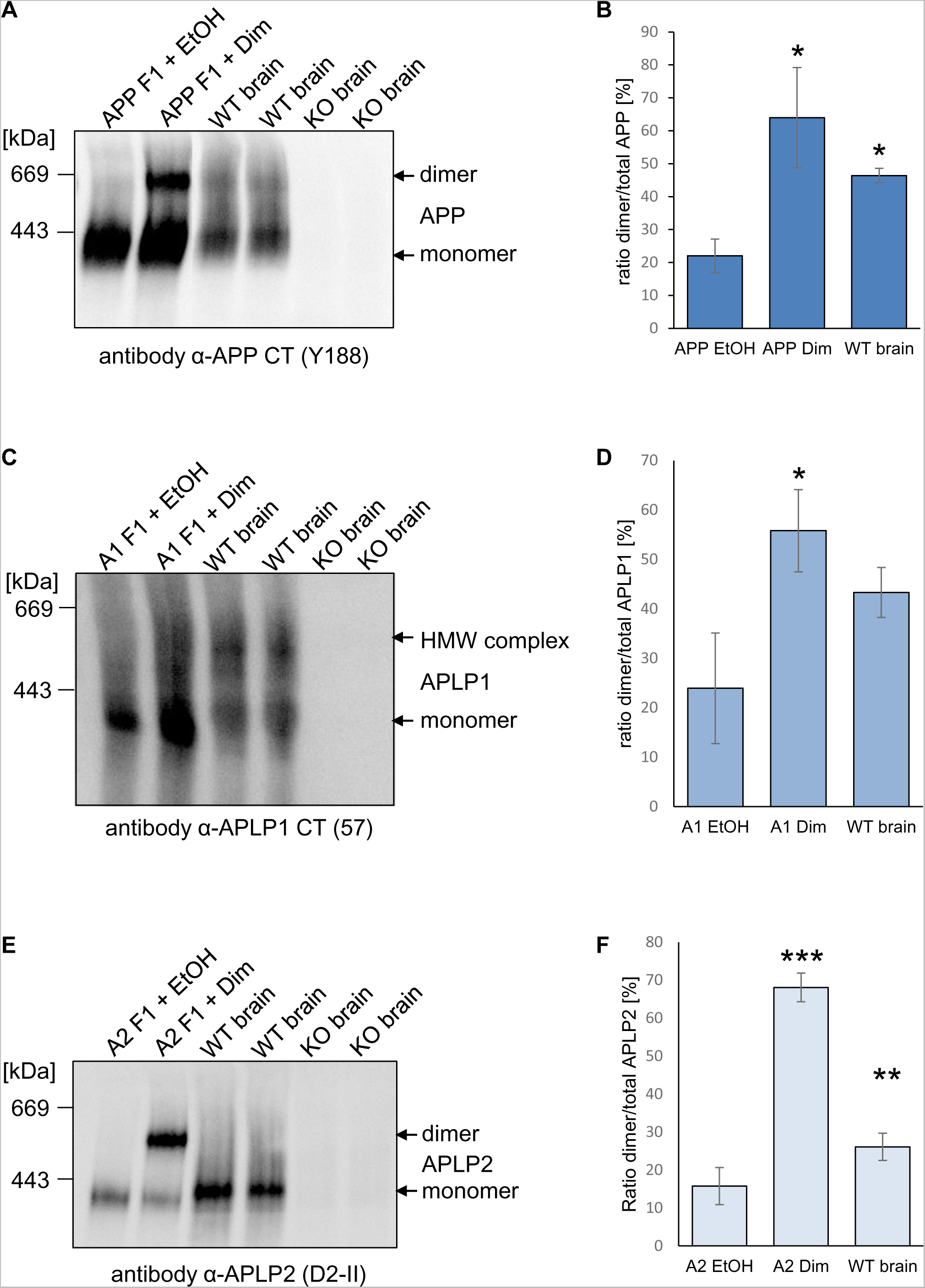
The extent of APP, APLP1, APLP2 dimers in N2a cells versus mouse brains. **(A)** APP F1 was heterologously expressed in N2a cells and treated either with 100 nM AP20187 over night to induce dimerization of APP or with the same volume of the vehicle ethanol as a control. One year old WT mouse cortices and APP KO mouse cortices were homogenized, and all samples were prepared under semi-denaturing conditions for analysis on Blue Native gels. Western blot detection of APP followed with the primary α-APP C-terminal antibody Y188. **(B)** Quantification of data shown in A. Bars represent mean values ± SEM of the ratio dimer/total of FKBP-rapamycin induced APP dimer in N2a cells and endogenous APP dimer in mouse brains. *n* = 3. **(C)** APLP1 F1 was heterologously expressed in N2a cells and treated either with 100 nM AP20187 over night to induce dimerization of APLP1 or with the same volume of the vehicle ethanol as a control. In parallel, one year old WT mouse cortices and APLP1 KO mouse cortices were analyzed, and all Blue Native gel samples were prepared under semi-denaturing conditions. Western blot detection of APLP1 followed with the primary α-APLP1 antibody 57. **(D)** Quantification of data shown in C. Bars represent mean values ± SEM of the ratio dimer/total of FKBP-rapamycin induced APLP1 dimer in N2a cells and endogenous APLP1 dimer in mouse brains. *n* = 3. **(E)** APLP2 F1 was heterologously expressed in N2a cells and treated either with 100 nM AP20187 over night to induce dimerization of APLP2 or with the same volume of the vehicle ethanol as a control. Those samples were compared to one year old WT mouse cortices and APLP2 KO mouse cortices. Analysis on Blue Native gels was followed by Western blot detection of APLP2 with the primary α-APLP2 antibody D2-II. **(F)** Quantification of data shown in E. Bars represent mean values ± SEM of the ratio dimer/total of FKBP-rapamycin induced APLP2 dimer in N2a cells and endogenous APLP2 dimer in mouse brains. *n* = 3.

As a next step, we wanted to analyze complex formation of the APP family members in cortical mouse brains in more detail. Therefore, we were examining dimer formation at the different developmental time points, E14, E17, P1, P3, P6, P10; P12, and P30 **(Figure 6A, C, E)**. The corresponding KO brains were used as a control for specificity of the antibodies. In addition, a longer period of APP, APLP1, and APLP2 complex formation was investigated in 1 month, 6 months, and 12-months-old cortical mouse brains **(Figure 6B, D, F)**.

**Figure 6:**
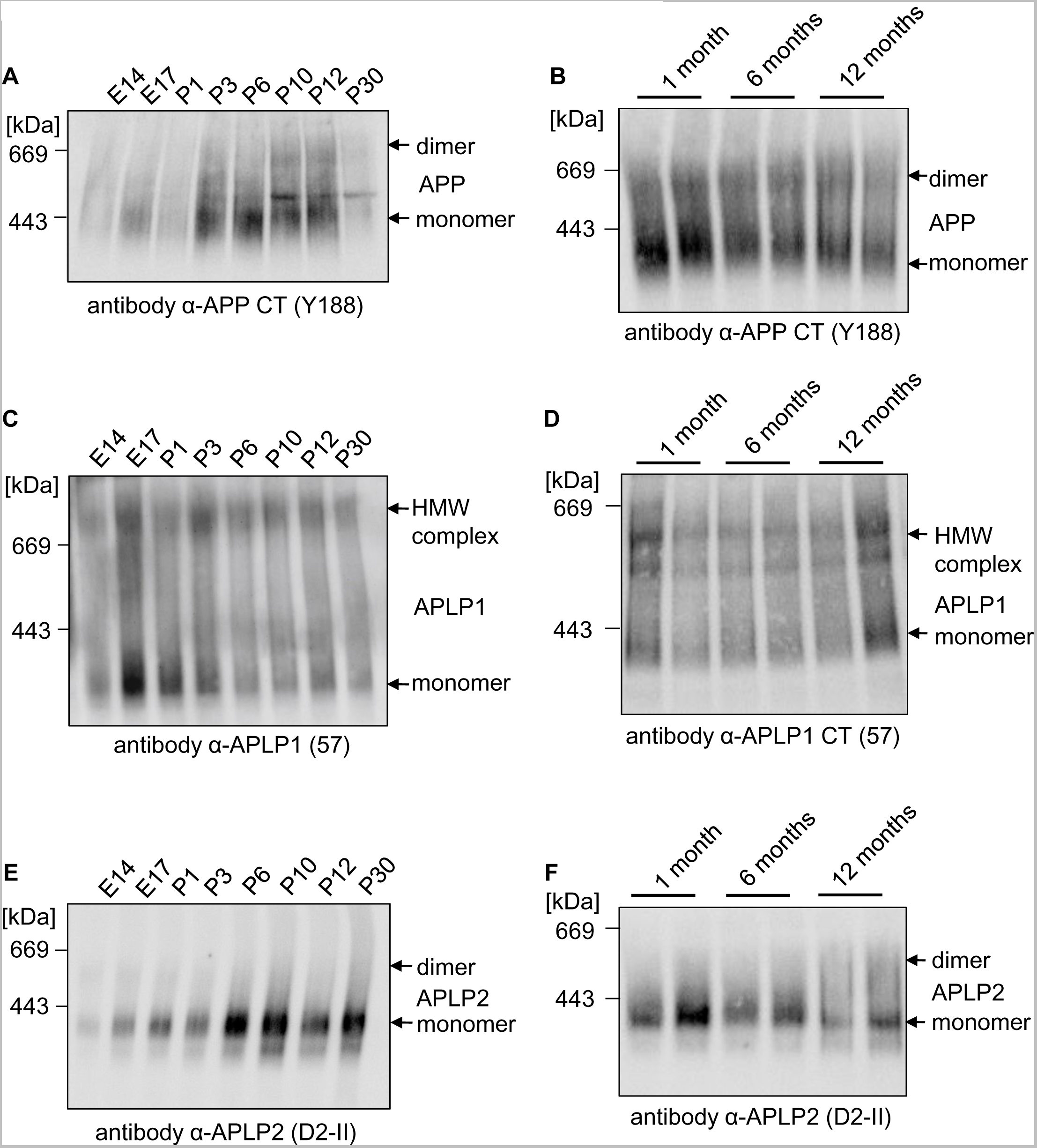
Dimer amount of the APP gene family members differs during development. **(A-F)** Expression analysis of dimers of the APP gene family members in mouse cortices during development. Mouse cortices of developmental stages E14, E17, P1, P3, P6, P10, P12, P30, 1 month, 6 months and 12 months were lysed. All samples were prepared under semi-denaturing conditions for analysis on Blue Native gels. Western blot detection of APP followed with the primary α-APP C-terminal antibody Y188 **(A, B)**, of APLP1 with the primary α-APLP1 antibody 57 **(C, D)** and APLP2 with the primary α-APLP2 antibody D2-II **(E, F)**.

APP was visualized with the KO verified specific antibody α-Y188 and increasing APP complex formation was detected with little dimer formation starting E14 and the peak at P10/P12 **(Figure 6A)**. APP dimer formation was unchanged between 1 month and 12 months old mice and constantly ∼49% **(Figure 6B)**. In contrast, formation of APLP1 high molecular weight complexes was unchanged between E14 and P30 and between 1 month and 12-months-old mice and constantly high **(Figure 6C, D)**.

Interestingly, APLP2 dimerization in cortical mouse brain was strongest E14 and continuously decreasing until P30 **(Figure 6E)**. The signal remained constantly low in cortical mouse brains between 1 month and 12 months of age **(Figure 6F)**.

Taken together, APP dimerization in cortical mouse brains peaked at P10/P12, the period of synapse formation and remained high at least until 12 months of age while APLP2 showed the opposite phenomenon with the highest amount of APLP2 dimers at E14, slowly decreasing and remaining constantly low until the age of one year. In contrast, APLP1 high molecular complex formation seems to be constant during development and during adulthood.

Next, we wanted to address the question why dimerization of APP is strongly increased in mouse brains compared to cell culture cells. For APP, we were analyzing so far splice form APP695 with the APP F1 construct. In general, eight different splice forms are known for APP whereby APP695, APP751, and APP770 are the three major isoforms being expressed, thereof APP695 primarily in neurons (Sandbrink et al., 1996b). Therefore, we were testing if differences regarding the degree of APP dimerization are splice form dependent. We generated C-terminally HA tagged constructs of APP695, APP751, and APP770 and verified the correct sizes of these proteins after SDS gel analysis and Western blot detection with an α-HA antibody **(Figure 7A)**. Next, we compared APP695 CT HA, APP751 CT HA, and APP770 CT HA with the inducible APP695 F1 dimerization system via BN gel analysis. Western blot detection with an α-HA antibody revealed that APP dimer formation was comparable between the different APP splice forms suggesting that increased APP dimer formation in cortical mouse brains is independent of splice variants **(Figure 7B)**. Next, we examined dimer formation of the APP family members in mouse cortical astrocytes, endogenously in mouse N2a cells and primary cortical neuronal cultures at an early stage DIV5 and a later stage which includes synapse formation (DIV19) **(Figure 8)**. Firstly, we were analyzing dimerization level of APP. APP dimers in cortical WT mouse brains showed again a percentage of ∼45% while there were only very little APP dimers present in primary astrocytes **(Figure 8A)** and primary cortical neuronal cultures (DIV5), but they were increased at DIV19 **(Figure 8B)**. APLP1 high molecular weight complexes in cortical WT mouse brains showed again a signal between 200 and 800 kDa, while there was no specific signal present for APLP1 in samples of cortical astrocytes, which was expected for the neuronal protein APLP1 (Lorent et al., 1995) and a very low signal in mouse neuronal N2a cells **(Figure 8C)**. APLP1 high molecular weight complexes between 200 and 800 kDa were also present in primary cortical neuronal cultures DIV5 and DIV19 but did not reveal any differences **(Figure 8D)**. APLP2 dimers in cortical WT mouse brains showed again a low percentage while there were surprisingly more APLP2 dimers present in primary astrocytes and mouse neuronal N2a cells **(Figure 8E)**. Primary cortical neuronal cultures contained also only a very low amount of APLP2 dimers at DIV5 and DIV19 **(Figure 8F)**. Taken together, APP dimerization seems to be increased in primary neuronal cultures at the time point of synapse formation and APLP2 dimerization is increased in primary cultures of cortical astrocytes while no modulation was found for APLP1 in these different culture systems.

**Figure 7:**
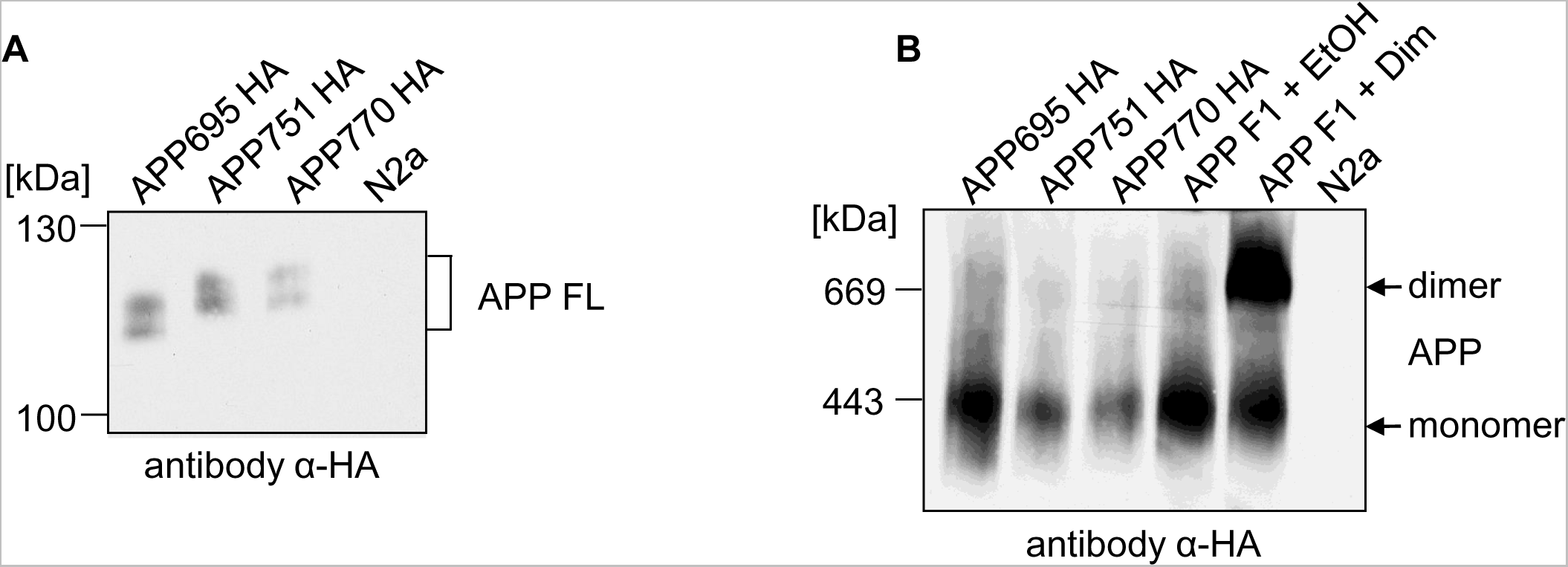
The dimer amount is independent of different APP splice variants. **(A)** APP695 HA, APP751 HA and APP770 HA were heterologously expressed in N2a cells. Non-transfected N2a cells served as a negative control. Equal amounts of cell lysates were separated on SDS-Gels and Western blot detection followed with an anti-HA antibody. **(B)** APP695 HA, APP751 HA and APP770 HA or APP F1 were heterologously expressed in N2a cells. To show APP dimers, APP F1 expressing cells were treated with 100 nM of the rapamycin analogue AP20187 overnight to induce dimerization of APP. Incubation with the solvent of AP20187, ethanol, served as a negative control. The samples were analyzed via Blue Native PAGE and Western blot analysis followed with an α-HA antibody.

**Figure 8:**
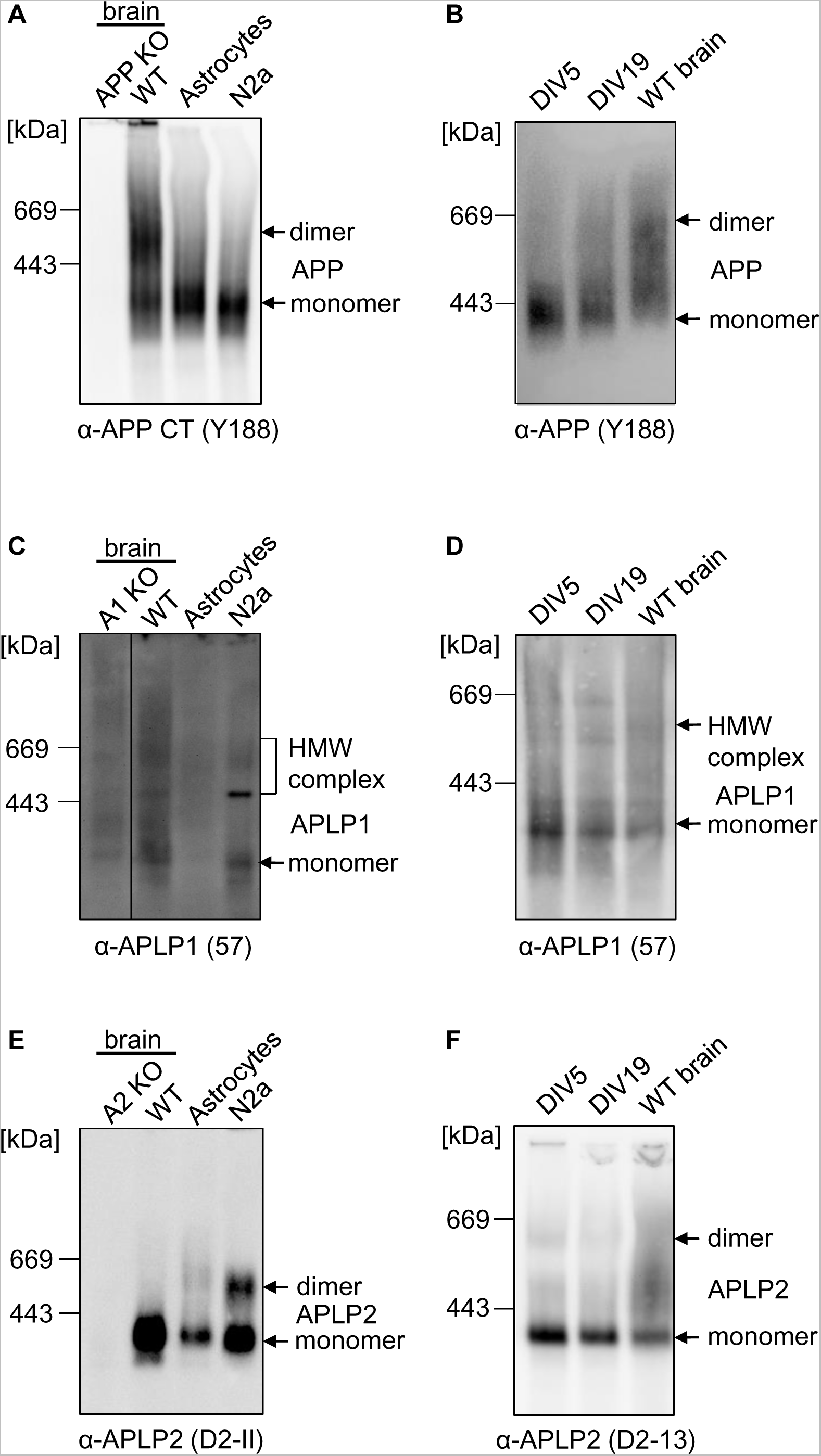
Higher amount of APP/APLP1/APLP2 dimerization is not visible in primary astrocytes, N2a cells or primary cortical neurons (DIV5). **(A/C/E)** One year old WT mouse cortices and APP KO mouse cortices were homogenized and compared with the lysate of astrocytes and N2a cells. All samples were prepared under semi-denaturing conditions for analysis on Blue Native gels. Western blot detection of APP followed with the primary α-APP C-terminal antibody Y188 **(A)**, of APLP1 with the primary α-APLP1 antibody 57 **(C)** and APLP2 with the primary α-APLP2 antibody D2-II **(E). (B/D/F)** Mouse primary cortical neurons were cultivated for 5 or 19 days *in vitro* (DIV), lysed and compared to the lysate of one year old WT mouse cortices. All samples were prepared under semi-denaturing conditions for analysis on Blue Native gels. Western blot detection of APP followed with the primary α-APP C-terminal antibody Y188 **(B)**, of APLP1 with the primary α-APLP1 antibody 57 **(D)** and APLP2 with the primary α-APLP2 antibody D2-II **(F).**

Via BN gel analyses, it can’t be distinguished if the visualized dimers form due to *cis-* or *trans*-dimerization. Based on the result of an increased presence of APP dimers in neuronal cultures at the time point of synapse formation, we were wondering if the higher rate of APP dimers in mouse brains is based on *trans*-dimerization at the synapse. Therefore, we were analyzing mouse brain synaptosomes which contain the synaptic terminals via BN gel analyses for all APP family members **(Figure 9)**. Synaptosomal fractions were prepared using WT and the corresponding cortical KO brains and examined via SDS PAGE and Western blot analysis. Synaptophysin and PSD 95 antibodies were used as a control for enrichment of pre- and postsynaptic fractions of the synapses, respectively **(Figure 9A, C, E)**. BN gel analysis revealed an obvious increase in the amount of APP dimers in enriched synaptosomal fractions, postsynaptic density H (PSD H), compared to post nuclear fractions (PNF) which contain crude cell lysates of the brain **(Figure 9B)**. The corresponding KO brain fractions were used as a control for specificity of the antibody signal. For APLP1, no major differences between the PNF and the PSD H fraction were observed **(Figure 9C)** while for APLP2, similarly to APP, the signal of APLP2 dimers was increased in the PSD H fraction compared with the PNF fraction **(Figure 9F)**. Conclusively, APP and APLP2 dimerization seem to be increased in synaptosomal fractions, presumably based on *trans*-dimerization at the synapse.

**Figure 9:**
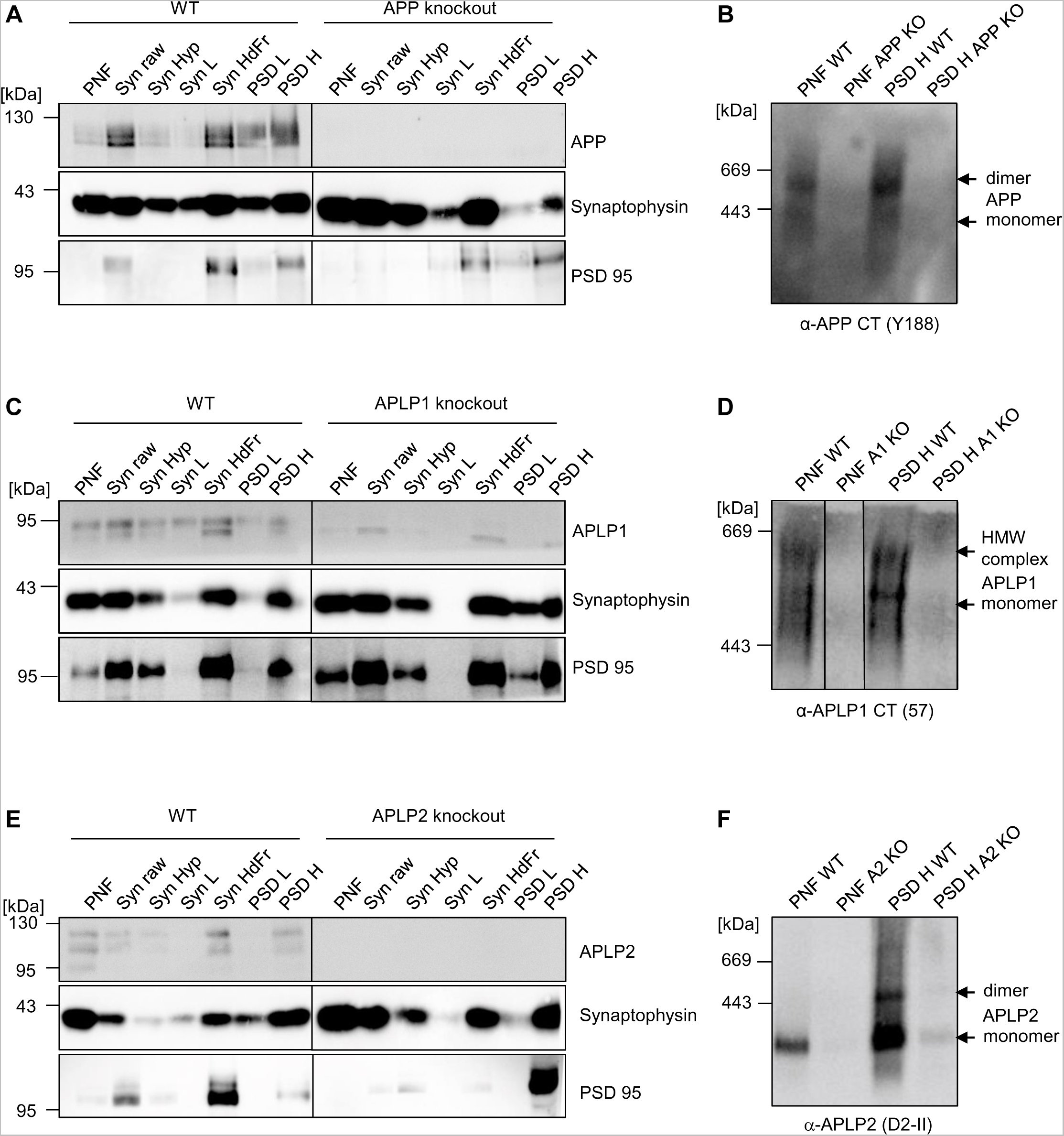
Expression analysis of APP, APLP1, and APLP2 dimers in mouse cortices in synaptosomal preparations. **(A/C/E)** WT mouse (C57BL/6) brain homogenates were sub fractionated by differential centrifugation steps and analyzed by Western blot with the primary α-APP C-terminal antibody Y188, α-APLP1 antibody 57, and α-APLP2 D2-II antibody. α-Synaptophysin and α-PSD-95 antibodies, respectively, were used as presynaptic and postsynaptic markers. To confirm specificity of the signals, synaptosome preparations were also performed with APP, APLP1, and APLP2 KO mouse brains. **(B/D/F)** Preparation of the post nuclear fraction (PNF) and the concentrated synaptosomes (PSD H) under semi-denaturing conditions for analysis of dimers on Blue Native gels. Western blot detection of APP followed with the primary α-APP C-terminal antibody Y188 **(B)**, of APLP1 with the primary α-APLP1 antibody 57 **(D)** and APLP2 with the primary α-APLP2 antibody D2-II **(F)**.

We next examined if familial AD (FAD) mutations show an impact on APP dimer formation. Therefore, we analyzed cortical mouse brains of J20 mice, which are transgenic mice encoding alternatively spliced minigenes of hAPP695, hAPP751, and hAPP770 containing the APP FAD mutations Swedish (K670N/M671L) and Indiana (V717F) under a neuronal PDGF-β promotor (Games et al., 1995; Mucke et al., 2000a; Rockenstein et al., 1995). J20 mice show the highest APP expression in the neocortex and hippocampus (Hong et al., 2016) and amyloid plaques are observed at 5-7 months in the dentate gyrus and neocortex in this AD mouse model (Hong et al., 2016; Mucke et al., 2000b). Conclusively, we investigated J20 cortical mouse brains for the presence of APP dimers at 4 months, before the onset of plaque formation and at 12 months when a high plaque load has developed. Non-transgenic littermates were used as a control. For the age of four months, we observed 58.14% ±1.25% APP dimers in J20 mice and 61.31% ±1.04% APP dimers in non-transgenic mice and for the age of 12 months 58.00% ±1.3% APP dimers in J20 mice and 54.75% ±1.8% APP dimers in non-transgenic mice. This indicates that there was no significant difference in the amount of dimer formation between murine APP and murine APP containing a humanized APP sequence including an FAD mutation at the β-secretase cleavage site and one at the γ-secretase cleavage site, also between two different age groups **(Figure 10 A-D)**. The result of increased APP dimer formation in the J20 mouse line implies that the observed APP dimers most likely result from all three major splice forms of APP (APP695, APP751, and APP770). Furthermore, the APP FAD mutations APP Swedish and APP Indiana do not seem to affect APP dimerization in mouse brains.

**Figure 10:**
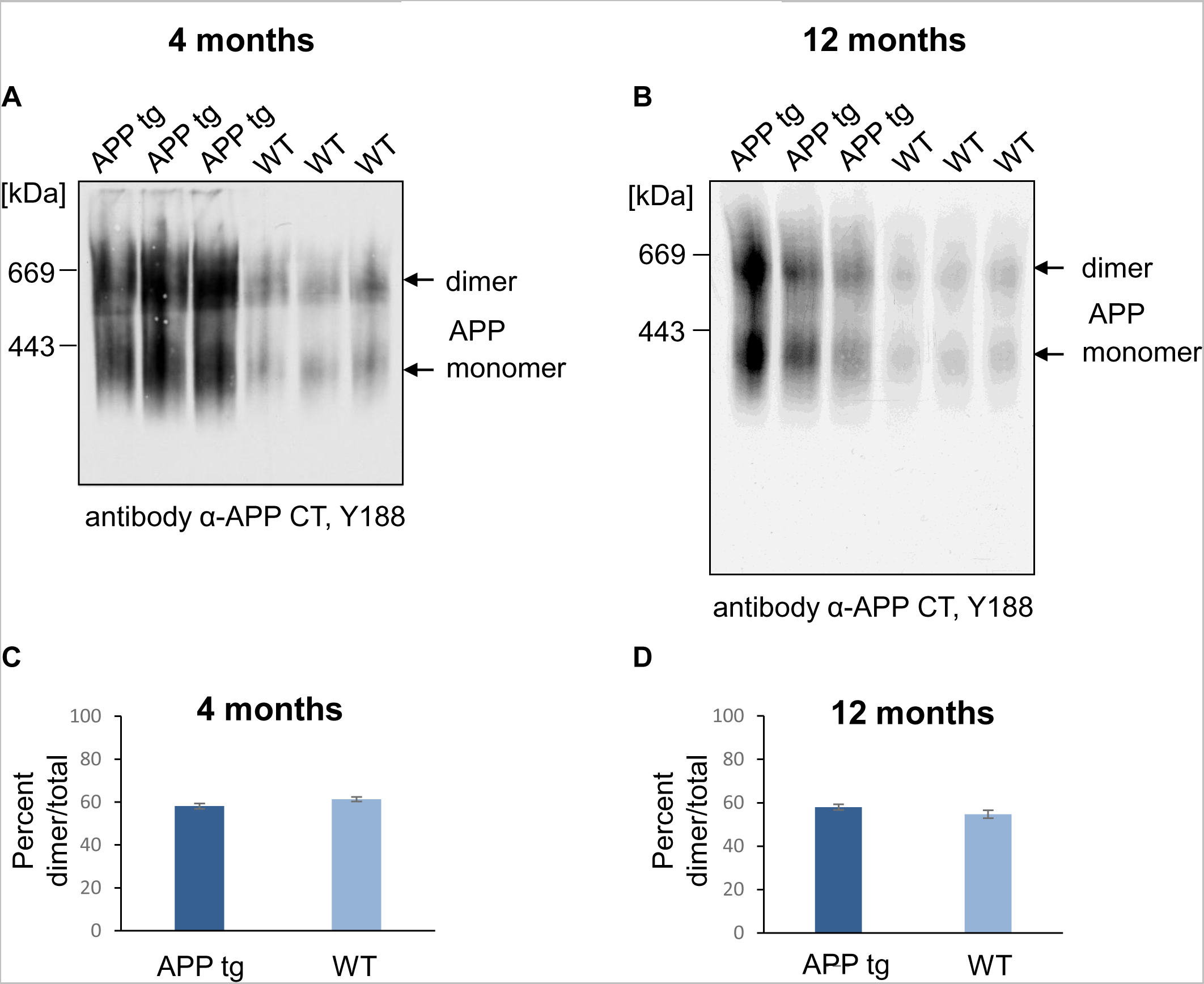
APP transgenic and WT mice show comparable amounts of APP dimer in the brain. **(A/B)** Mouse cortices of WT and of APP J20 transgenic mice at the age of 4 and 12 months. All samples were prepared under semi-denaturing conditions for analysis on Blue Native gels. Western blot detection of APP followed with the primary α-APP C-terminal antibody Y188. **(C/D)** Quantification of the data shown in A and B. Bars represent mean values ± SEM of dimer ratio to the total amount of the signal in %. *n* = 3, unpaired Student’s *t* test \**p* < 0.05, \*\**p* < 0.01, \*\*\**p* < 0.001.

To address the question if APP dimerization plays a role during AD, age matched, and area matched human AD brain samples of the middle frontal gyrus from sporadic AD patients were analyzed via BN gel analysis **(Figure 11)**. Quantification revealed a significant decrease in the amount of APP dimers in brains of AD patients, suggesting that APP dimerization gets impaired during the course of AD.

**Figure 11:**
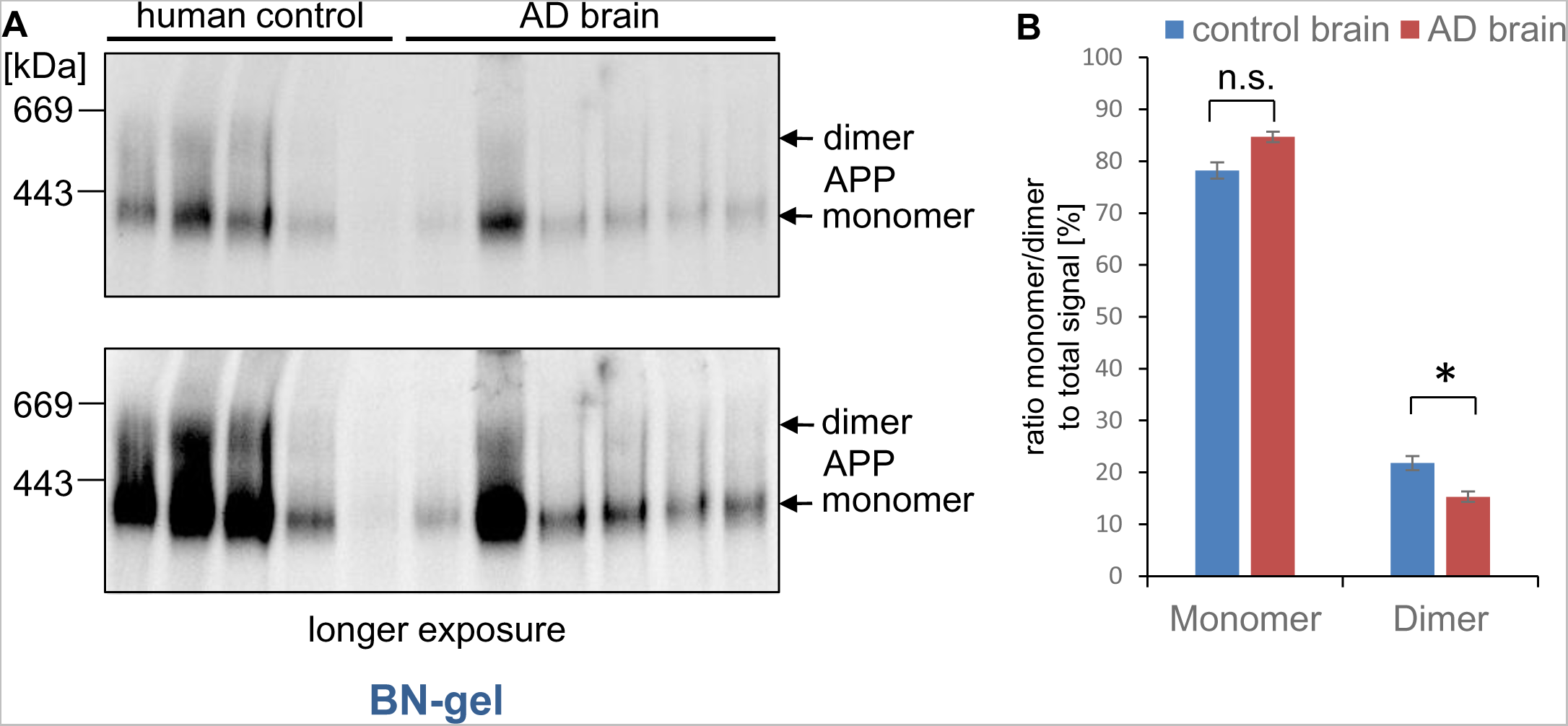
Alzheimer’s disease patients show decreased amounts of APP dimers in the brain. **(A)** Samples of the middle frontal gyrus of AD patients and human controls, age matched, and area matched, were lysed under semi-denaturing conditions for analysis on Blue Native gels. Western blot detection of APP followed with the primary anti-APP C-terminal antibody Y188. **(B)** Quantification of data shown in A. Bars represent mean values ± SEM of monomer/dimer ratio to the total amount of the signal in %. *n* = 3, unpaired Student’s *t* test \**p* < 0.05, \*\**p* < 0.01, \*\*\**p* < 0.001.

## DISCUSSION

### Comparison of APP, APLP1, and APLP2 complexes

APP, APLP1, and APLP2 can form dimers in *cis-* as well as in *trans-*orientation predominantly via the E1 domain in the extracellular area (Baumkotter et al., 2012; Kaden et al., 2009; Soba et al., 2005). On the other hand, the E2 domain has been found to dimerize as an antiparallel dimer suggesting that this domain might be involved in *trans*-dimerization of APP as well (Wang and Ha, 2004).

The degree of APP dimerization in cell culture cells has already been investigated using BN gel analyses and an APP-FKBP fusion protein (Eggert et al., 2009). With this system, after addition of the dimerization agent AP20187, ∼66% APP dimers could be induced in N2a cells, while only about 22% APP dimers were present in the control **(Figure 1, 5)** (Eggert et al., 2009). This suggests that approximately 22% of APP molecules are dimerized in cells, cultivated *in vitro*, which is in line with other independent reports (Sato et al., 2007; So et al., 2013). Furthermore, cross-linking studies in human SH-SY5Y cells corroborate the low proportion of APP *cis*-dimers in cell culture cells (Scheuermann et al., 2001; Soba et al., 2005) and also FRET experiments indicated APP homo-dimerization to a degree of 11% in HEK cells (Kaden et al., 2009; Munter et al., 2007).

Together, our data suggest that APP dimers represent only a minor proportion of total APP in cells, cultivated *in vitro*. The amount of *trans*-dimers is assumed to be even lower under these conditions. Cell surface localization of proteins is the prerequisite for their interaction in *trans*-orientation. Therefore, studies showing a low amount of APP at the cell surface suggest that only a low proportion of APP dimerizes in *trans*-orientation (Eggert et al., 2018a). The pattern of complexes on BN-gels obtained from transiently transfected N2a cells was quite comparable between APP and APLP2 showing the majority of the signal to be in the monomeric form **(Figure 1, 2)**. In contrast, APLP1 revealed a pattern of steady complex formation between 200 and 800 kDa, involving also dimerized complexes **(Figure 1, 2)**. The reason for the pattern of diverse high molecular weight complexes might be based on heparin or zinc induced APLP1 dimerization (August et al., 2019; Dahms et al., 2015; Dunsing et al., 2017; Mayer et al., 2014; Xue et al., 2011), differently glycosylated complexes (Paliga et al., 1997; Schilling et al., 2017), or processing by the transmembrane serine protease Matriptase, which cleaves APLP1 within the E1 domain and thereby negatively impacts APLP1 homodimerization (Lanchec et al., 2020).

Our results fit to the fact that APP and APLP2 share some characteristics in contrast to the family member APLP1, which has some unique features. APLP1 is exclusively present in neurons (Lorent et al., 1995) while APP and APLP2 are expressed ubiquitously (Wasco et al., 1993). For APLP1, only one isoform has been reported (Paliga et al., 1997; Wasco et al., 1992), whereas alternative splicing leads to four mRNA variants for APLP2 (Sandbrink et al., 1996a) and eight splice variants for APP (Sandbrink et al., 1994). Also, a higher presence for APLP1 compared with APP or APLP2 at the cell surface has been reported via cell surface biotinylation (Schilling et al., 2017) and ICC (Kaden et al., 2009) which we confirmed in our study **(Figure 2)**. Additionally, we could demonstrate that deleting the C-terminus of APP or APLP2 including their internalization motifs, shifts their presence to the cell surface, resulting in a higher amount of *trans*-dimers in eukaryotic cell lines in the case of APP **(Figure 2A, B)**. Regarding *trans*-dimerization of the APP family members, we already showed via an *in vitro* assay that APLP1 has the strongest *trans*-interacting properties and APP revealed the lowest degree of *trans*-dimerization (Schilling et al., 2017), which is in line with APLP1 having the strongest impact of the APP family members on presynaptic differentiation, presumably via *trans*-interaction with APLP1 in the presence of APLP2 (Schilling et al., 2017).

Together, we observed in heterologously expressing N2a cells, a relatively low amount of APLP2 and APP homotypic dimers while APLP1 showed a clearly higher degree of complex formation. Notably, other studies, using FRET analysis of transiently transfected HEK cells expressing APLP1 and APLP2 homo-dimers, reported that about 16% of total APLP1 and APLP2 are dimerized (Kaden et al., 2009). However, FRET analysis only allows detection of two fusion proteins (e.g. APLP1 fused to CFP and YFP) in close proximity, but does not test for direct binding. The low amount of detected APLP1 dimerization via FRET in contrast to our results might be based on conformational differences of the APLP1 fusion proteins or might be affected by the length of the linker region. Our co-immunoprecipitation data and BN gel analysis, indicate a strong interaction of APLP1 in a direct or indirect fashion **(Figure 1, 2, 3, 5)**.

### Diversity of APP, APLP1, and APLP2 complex formation in mouse brains

Highly interesting, we observed for APP a clearly elevated amount of high molecular weight complexes in brain homogenates compared to cell culture cells **(Figure 5)**. In contrast, the degree of APLP1 and APLP2 dimerization, although very different, did not vary between cell culture and mouse brain homogenates **(Figure 5)**. Also, developmental alterations in APP, APLP1, and APLP2 complex formation differed between the APP family members. APP complex formation was mostly increased when APLP2 complex formation was low, and vice versa **(Figure 6)**, suggesting that APP, APLP1, and APLP2 dimerization are underlying a regulation that depends on developmental time point and cell type **(Figure 8)**.

In this work, we were able to demonstrate that APP complex formation is increased in mouse cortices compared to cell culture cells from 22% to ∼49% **(Figure 5)**. The signals for APP monomers and dimers in mouse brain have been identified at ∼300 kDa and ∼600 kDa by comparison with dimerized and non-dimerized APP FKBP in N2a cells **(Figure 5)**. In contrast, APP monomers and APP dimers covalently induced by cysteine mutants APP L17C and APP K28C show an apparent molecular weight of ∼120 kDa and ∼240 kDa after separation in SDS gels (Eggert et al., 2018a) suggesting that APP is complexed with one or more binding partner, which is visible after BN-gel analysis.

The increased occurrence of APP dimers in mouse cortices compared to cell culture cells could be related to more *trans*-dimers being formed in intact tissue compared with cell culture cells. In cell culture, cell-cell contacts are built at higher confluency for a short period of time during cultivation and due to increased presence at the cell surface **(Figure 4)**. One reason for this might be APP - APP *trans*-interaction at the synapse. Consistently, APP was shown to localize at both, the pre- and postsynapse (Hoe et al., 2009; Schilling et al., 2017), which is the basic requirement for binding in *trans*-orientation at the synapse. Furthermore, synaptic *trans*-dimerization of APP proteins has been reported to promote synapse formation (Schilling et al., 2017; Soba et al., 2005; Stahl et al., 2014; Wang et al., 2009). Moreover, this assumption is supported by our data, showing that APP dimerization in mice has a peak at P10/P12, the period of synapse formation in mouse brains **(Figure 6)** and that APP dimers are increased in synaptosomes **(Figure 9)**.

Several factors have been described that influence APP dimer formation, such as Sortilin-Related Receptor Containing LDLR A Repeats (SorLA). SorLA is a protein that controls processing and intracellular transport of APP and impairs the formation of APP dimers in the brain (Schmidt et al., 2012). SorLA KO mice show significantly higher amounts of APP dimers with a simultaneously strong reduction in APP monomers (Schmidt et al., 2012) suggesting that SorLA promotes the formation of APP monomers and inhibits APP dimerization.

Furthermore, copper induces dimer formation of APP in both, *cis-* and *trans*-orientation (August et al., 2019; Baumkotter et al., 2014) and also Cholesterol lowering drugs seem to promote APP dimerization (Langness et al., 2021). Another factor inducing APP dimerization is the glycosaminoglycan heparin (Dahms et al., 2010; Faham et al., 1996). It has been reported that APP seems to be almost completely dimerized at the cell surface and involves the presence of heparan sulfate at the plasma membrane (Gralle et al., 2009). Surface localization of APP is a balance from its transport in the secretory path, its internalization and its processing (Eggert et al., 2018a; Eggert et al., 2020; Eggert et al., 2022; Eggert et al., 2018b). For example, reduced proteolysis of APP in processing deficient mutants leads to an increased surface localization of APP, resulting in elevated *trans*-dimerization and synapse formation (Stahl et al., 2014). APP internalization from the cell surface occurs predominantly by Clathrin mediated endocytosis via the NPTY motif in the C-terminus of APP (Eggert et al., 2020; Koo and Squazzo, 1994), but also via the basolateral Sorting sequence (BaSS) of APP which is the YTSI motif C-terminal of the transmembrane domain (Eggert et al., 2020; Lai et al., 1995).

APP, a type I transmembrane protein has the structure of a cell surface receptor (Kang et al., 1987) as also its homologues APLP1 and APLP2. It is discussed that localization of APP at the plasma membrane allows the interaction with other proteins, such as the APP homologous proteins via the E1 domain (Deyts et al., 2016). The APP ectodomain has been reported to adopt different conformations depending on cellular pH (Hoefgen et al., 2015). A pH-dependent interaction between the GFLD subdomain with the copper-binding domain (CuBD) within the E1 domain might lead to a closed conformation for APP in a more acidic compartment like endosomes, whereas it will adopt a more opened conformation at neutral pH 7.4, which is present at the cell surface (Hoefgen et al., 2015). Therefore, APP in its open conformation at the plasma membrane can expose the E1 domain to allow association with other receptors or ligands resulting in different physiological functions. This is consistent with noticeable high molecular weight complex formation of the APP family members after BN gel electrophoresis **(Figure 1, 2, 5)**. The APP protein has only a very short half-life time of about 45 minutes (Weidemann et al., 2002; Weidemann et al., 1989) implicating only a brief dwell time of APP at the plasma membrane. This indicates that *trans*-dimerization of APP, homophilic or heterophilic, is most likely not essential for stabilization of cell-cell interactions but will rather induce signal transduction processes. One interesting APP interaction partner is the protein N-cadherin, which is involved in initiation of synapse formation (Tamura et al., 1998) reported to increase APP dimerization in *cis*-orientation (Asada-Utsugi et al., 2011).

It was shown that *cis*-homo-dimerization of APP and/or the β-C-terminal fragment significantly reduced the amount of total Aβ (Eggert et al., 2009; Jung et al., 2014; Langness et al., 2021; Winkler et al., 2015). This suggests that more APP monomers lead to increased Aβ generation, which in turn should lead to more Aβ oligomers and ultimately more Aβ plaques. Thus, higher amounts of APP dimers might have a long-term positive effect in Alzheimer’s disease patients. In this work, it could be shown that APP in human cortex samples from non-demented controls is 21.8 ± 1.3 % as a dimer and 78.2 ± 1.3 % as a monomer **(Figure 11)**. Interestingly, the proportion of APP dimers is significantly reduced in samples from Alzheimer’s disease patients **(Figure 11)**. This could indicate that APP *trans*-dimerization is severely reduced in Alzheimer’s brains and that factors influencing this interaction play an important role in Alzheimer’s disease, since it is known that progressive Alzheimer’s dementia coincides with loss of synapse formation (Scheff et al., 2007). Thus, pharmacological approaches targeting APP oligomerization properties might open novel strategies for treatment of AD.

## Acknowledgements

We thank Luigina Hanke for excellent administrative assistance. We thank Alzheimer Forschung Initative e.V., DFG (KI 819/9-1) and BioComp (Forschungsinitiative Rheinland-Pfalz) for funding to SK and Alzheimer Forschung Initative e.V (19084) and BioComp to SE. SE was supported by funding of the TU Nachwuchsring. The authors declare no competing financial interests.

